# A Novel Homeostatic Mechanism Tunes PI(4,5)P_2_-dependent Signaling at the Plasma Membrane

**DOI:** 10.1101/2022.06.30.498262

**Authors:** Rachel C. Wills, Colleen P. Doyle, James P. Zewe, Jonathan Pacheco, Scott D. Hansen, Gerald R. V. Hammond

## Abstract

The lipid molecule phosphatidylinositol (4,5)-bisphosphate (PI(4,5)P_2_) controls all aspects of plasma membrane (PM) function in animal cells, from its selective permeability to the attachment of the cytoskeleton. Although disruption of PI(4,5)P_2_ is associated with a wide range of diseases, it remains unclear how cells sense and maintain PI(4,5)P_2_ levels to support various cell functions. Here, we show that the PIP4K family of enzymes that synthesize PI(4,5)P_2_ via a minor pathway, also function as sensors of tonic PI(4,5)P_2_ levels. PIP4Ks are recruited to the PM by elevated PI(4,5)P_2_ levels, where they inhibit the major PI(4,5)P_2_-synthesizing PIP5Ks. Perturbation of this simple homeostatic mechanism reveals differential sensitivity of PI(4,5)P_2_-dependent signaling to elevated PI(4,5)P_2_ levels. These findings reveal that a subset of PI(4,5)P_2_-driven functions may drive disease associated with disrupted PI(4,5)P_2_ homeostasis.

**One-Sentence Summary:** The enzyme PIP4K functions as both a sensor and negative regulator of PI(4,5)P_2_ synthesis by the closely related PIP5K enzymes, tuning the activity of numerous membrane functions.

## Introduction

The lipid molecule PI(4,5)P_2_ is a master regulator of animal cell plasma membranes (PMs). By recruiting or activating scores of membrane proteins, it controls transport of ions and solutes across the membrane [1,2], attaches the underlying cytoskeleton [3], regulates the traffic of proteinaceous cargo to and from the membrane [4], disseminates extracellular signals [2], and facilitates the entry, assembly and egress of bacterial and viral pathogens [2,5]. As a result, synthesis of PI(4,5)P_2_ is essential for life in mammals [6,7]. Nonetheless, genetic defects occur in humans that either increase or decrease PI(4,5)P_2_ levels, disrupting cellular physiology in unpredictable ways. These manifest in diseases ranging from cancer [8] to kidney disease [9] to dysentery [10]. Clearly, there is a central physiological imperative to tightly control PI(4,5)P_2_ levels for harmonious PM function. A detailed homeostatic mechanism that can sense and maintain PI(4,5)P_2_ levels has, however, proven elusive.

Most prior work in this area has focused on positive regulation of phosphatidylinositol 4-phosphate 5-kinases (PIP5Ks), the major enzymes responsible for PI(4,5)P_2_ synthesis (**fig. 1A**). These enzymes add a phosphate to the 5-OH of their substrate, PI4P [11–13]. Such positive regulation can be mediated by the small GTPases Arf6 [12,13] and Rac [14,15] or the PI(4,5)P_2_ metabolite phosphatidic acid [16]. In fact, PIP5Ks cooperatively bind to their product, PI(4,5)P_2_, which creates a positive feedback loop that enhances membrane localization and catalytic output [17]. However, we reasoned that maintaining tonic PI(4,5)P_2_ levels in the PM in the presence of abundant PI4P substrate [18,19] would demand negative feedback of PIP5Ks. This is especially apparent during lipid re-synthesis after phospholipase C (PLC) activation; PI(4,5)P_2_ levels plateau despite the fact that levels of the precursor lipid PI4P are still rising [20–22]. Potential mechanisms of PI(4,5)P_2_ downregulation include PIP5K autophosphorylation [23], as well as a futile cycle wherein PI(4,5)P_2_ lipids are dephosphorylated back to PI4P by inositol polyphosphate 5-phosphatase (INPP5) enzymes [24], although the specific INPP5 family member(s) responsible for this constitutive activity have not been defined. Finally, PIP5K inhibition by the related phosphatidylinositol 5-phosphate 4-kinases (PIP4Ks), which produce PI(4,5)P_2_ from much less abundant PI5P substrate, has been reported [25]. However, how this downregulation of PIP5K activity by the PIP4Ks is regulated to maintain PI(4,5)P_2_ homeostasis has not been defined.

**Fig. 1.**
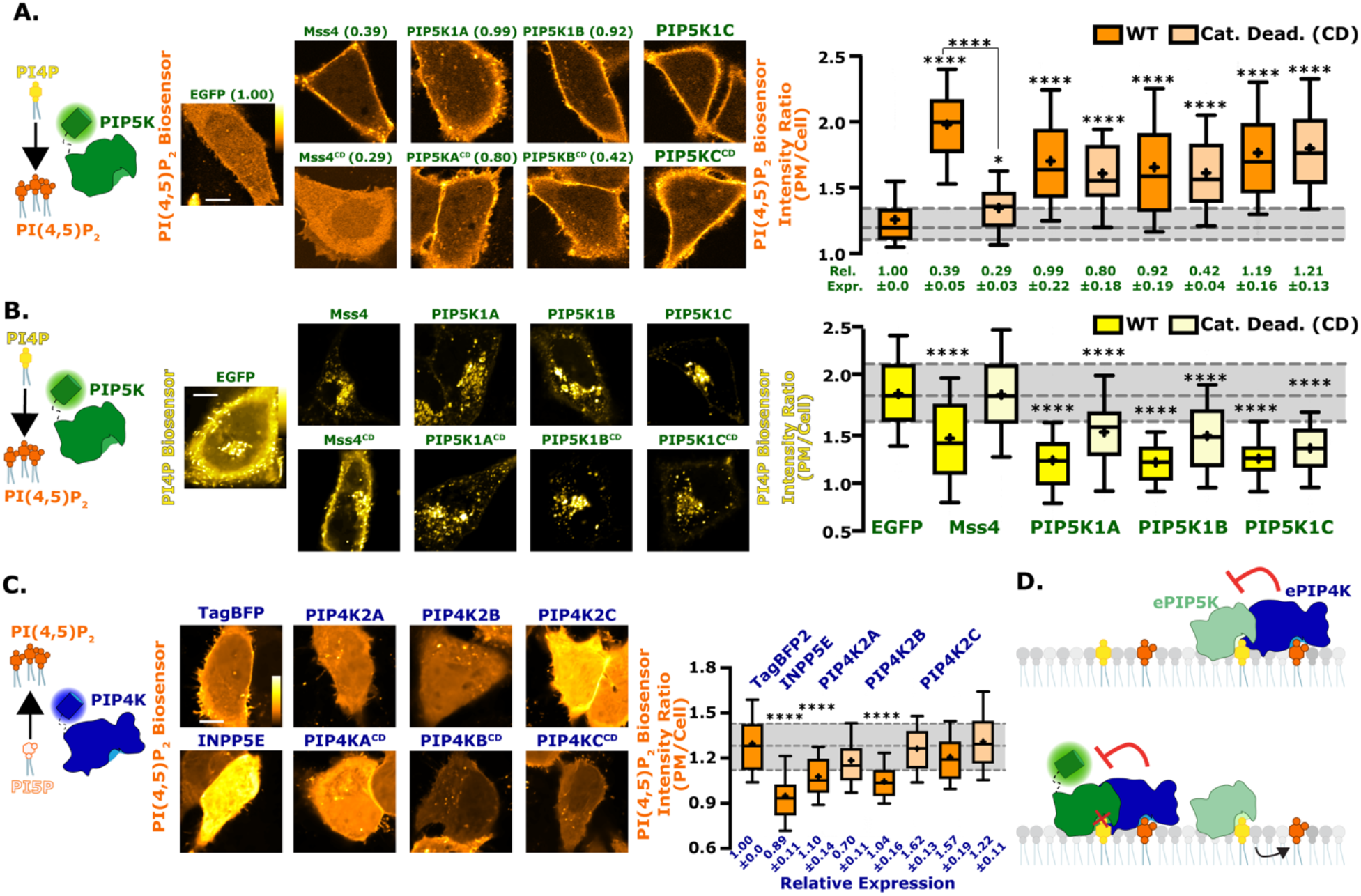
Reciprocal regulation of PM PI(4,5)P_2_ levels by PIP5K and PIP4K. (**A**) PIP5Ks increase PM PI(4,5)P_2_ independently of catalytic activity. Cartoon denotes the catalytic activity of PIP5K. Images show equatorial confocal sections of HeLa cells expressing the low affinity TubbyC^R332H^ PI(4,5)P_2_ sensor (orange), co-transfected with EGFP-tagged catalytically active or dead PIP5K constructs (yeast Mss4 or mammalian A, B, or C paralogs), or EGFP alone as control, scale bar is 10 µm. Increased PI(4,5)P_2_ is apparent from higher TubbyC^R332H^ fluorescence in the PM. Box and whisker plots show the mean fluorescence intensity ratio (PM/Cell) of the PI(4,5)P_2_ sensor from > 90 cells imaged across at least three independent experiments (boxes displaying median and interquartile range, whiskers representing 10-90% of data and “+” represents mean; *P < 0.05; **P < 0.01; ***P < 0.001; ****P < 0.0001). Expression of EGFP-tagged proteins relative to the EGFP control is indicated below box and whisker plots based on raw fluorescence intensity; data are means ± s.e. (**B**) PIP5Ks reciprocally decrease PM PI4P independently of catalytic activity. Cartoon denotes the catalytic activity of PIP5K. Images show equatorial confocal sections of HeLa cells expressing the high affinity P4Mx2 PI4P sensor (yellow), co-transfected with EGFP-tagged catalytically active or dead PIP5K constructs (yeast Mss4 or mammalian A, B, or C paralogs), or EGFP alone as control, scale bar is 10 µm. Decreased PI4P is apparent from loss of P4Mx2 fluorescence at the PM. Box and whiskers from 88-90 cells imaged across at least three independent experiments as in A. (**C**) PIP4Ks decrease PM PI(4,5)P_2_ independently of catalytic activity. Cartoon shows the catalytic activity of PIP4K. Images show PI(4,5)P_2_ sensor in HeLa cells as in A, co-transfected with different PIP4K paralogs (A, B, C), catalytically dead PIP4K2A or a PI(4,5)P_2_ 5-phosphatase(pptase) (INPP5E), scale bar is 10 µm. Box and whiskers from > 90 cells imaged across at least three independent experiments as in A. Expression of TagBFP2-tagged proteins relative to the TagBFP_2_ control is indicated below box and whisker plots based on raw fluorescence intensity; data are means ± s.e. (**D**) Proposed inhibition of ePIP5K (endogenous PIP5K) by ePIP4K. With the overexpression of a fluorescently tagged version of PIP5K, regardless of catalytic activity, ePIP4K is sequestered. This relieves endogenous PIP5K from inhibition, increasing PI(4,5)P_2_ levels.

A common feature missing from effectors that downregulate PI(4,5)P_2_ synthesis is the identity of sensors that detect changing PI(4,5)P_2_ levels and modulate these effectors appropriately. Without knowledge of such a mechanism, how cells accomplish effective PI(4,5)P_2_ homeostasis and thereby maintain harmonious PM function has been a mystery. In this paper, we demonstrate that the PIP4K family of enzymes act as low affinity PI(4,5)P_2_ sensors, monitoring tonic PI(4,5)P_2_ levels and constraining PIP5K activity when levels of the lipid rise too high. Modulation of this homeostatic mechanism reveal unprecedented differences in the sensitivity of PI(4,5)P_2_-dependent signaling to resting PI(4,5)P_2_ levels.

## Results

### PIP5Ks are inhibited by PIP4Ks

This study was motivated by some initially perplexing results we obtained when monitoring PM PI(4,5)P_2_ levels with the low-affinity biosensor, Tubby_c_^R332H^ [26], together with PI4P levels using the high-affinity biosensor, P4Mx2 [19]: PI(4,5)P_2_ levels are expected to increase at the expense of PM PI4P levels when over-expressing any of the three paralogs of human PIP5K (A-C) or the single homolog from the budding yeast, *Saccharomyces cerevisiae* (Mss4). Indeed, this was precisely what we observed (**fig. 1A and B**, statistics reported in **tables 1 and 2**). What perplexed us was that catalytic activity of the human enzymes is dispensable for increased PI(4,5)P_2_ (**fig. 1A**) and depleted PM PI4P (**fig. 1B**). Catalytic activity is essential for yeast PIP5K, however (**fig. 1A and B**). Notably, expression of the catalytically inactive mutants was usually somewhat less strong compared to the wild-type enzymes, yet effects on PI(4,5)P_2_ levels were similar (**fig. 1A**).

**Table 1:** P values from Kruskal-Wallis test with Dunn’s correction for multiple comparisons for Tubbyc^R332H^ (PI(4,5)P_2_) biosensor data presented in Figure 1A. Significant variation was observed among groups by one-way ANOVA (KW statistic = 5089, P <0.0001). Significant results are highlighted in bold and n values are displayed.

| Construct: | EGFP<br>(n=2294) | Mss4 | Mss4<sup>CD</sup> | PIP5K1A | PIP5K1A<sup>CD</sup> | PIP5K1B | PIP5K1B<sup>CD</sup> | PIP5K1C |
| --- | --- | --- | --- | --- | --- | --- | --- | --- |
| Mss4 (n=90) | **<0.0001** |  |  |  |  |  |  |  |
| Mss4<sup>CD</sup><br>(n=90) | **0.0324** | **<0.0001** |  |  |  |  |  |  |
| PIP5K1A<br>(n=299) | **<0.0001** | **<0.0001** | **<0.0001** |  |  |  |  |  |
| PIP5K1A<sup>CD</sup><br>(n=90) | **<0.0001** | **<0.0001** | **<0.0001** | >0.9999 |  |  |  |  |
| PIP5K1B<br>(n=180) | **<0.0001** | **<0.0001** | **<0.0001** | >0.9999 | >0.9999 |  |  |  |
| PIP5K1B<sup>CD</sup><br>(n=90) | **<0.0001** | **<0.0001** | **0.0001** | >0.9999 | >0.9999 | >0.9999 |  |  |
| PIP5K1C<br>(n=1729) | **<0.0001** | **0.0003** | **<0.0001** | >0.9999 | 0.4741 | **0.0094** | 0.2673 |  |
| PIP5K1C<sup>CD</sup><br>(n=384) | **<0.0001** | **0.0347** | **<0.0001** | 0.3338 | 0.0743 | **0.001** | **0.0396** | >0.9999 |

**Table 2:** P values from Kruskal-Wallis test with Dunn’s correction for multiple comparisons for P4Mx2 (PI4P) biosensor data presented in Figure 1B. Significant variation was observed among groups by one-way ANOVA (KW statistic = 231.2, P <0.0001). Significant results are highlighted in bold and n values are displayed.

| Construct | EGFP<br>(n=90) | Mss4 | Mss4<sup>CD</sup> | PIP5K1A | PIP5K1A<sup>CD</sup> | PIP5K1B | PIP5K1B<sup>CD</sup> | PIP5K1C |
| --- | --- | --- | --- | --- | --- | --- | --- | --- |
| Mss4 (n=90) | **<0.0001** |  |  |  |  |  |  |  |
| Mss4<sup>CD</sup><br>(n=90) | >0.9999 | **<0.0001** |  |  |  |  |  |  |
| PIP5K1A<br>(n=90) | **<0.0001** | **0.0325** | **<0.0001** |  |  |  |  |  |
| PIP5K1A<sup>CD</sup><br>(n=90) | **<0.0001** | >0.9999 | **0.0004** | **<0.0001** |  |  |  |  |
| PIP5K1B<br>(n=90) | **<0.0001** | **0.0108** | **<0.0001** | >0.9999 | **<0.0001** |  |  |  |
| PIP5K1B<sup>CD</sup><br>(n=90) | **<0.0001** | >0.9999 | **<0.0001** | **0.0008** | >0.9999 | **0.0002** |  |  |
| PIP5K1C<br>(n=90) | **<0.0001** | 0.1166 | **<0.0001** | >0.9999 | **0.0001** | >0.9999 | **0.004** |  |
| PIP5K1C<sup>CD</sup><br>(n=90) | **<0.0001** | >0.9999 | **<0.0001** | 0.6371 | 0.3208 | 0.2742 | >0.9999 | >0.9999 |

Conversely, over-expression of PIP4K enzymes, which also make PI(4,5)P_2_ but from PI5P substrate, would be expected to elevate PI(4,5)P_2_ levels slightly. Yet, we found that PIP4K2A and PIP4K2B actually decreased PM PI(4,5)P_2_ levels, with a ranked order PIP4K2A > PIP4K2B > PIP4K2C (**fig. 1C**, statistics in **table 3**). For PIP4K2A at least, this occurred even when expressing a catalytically inactive mutant. Again, differences in expression level between paralogs do not explain differences in activity, since all achieved comparable expression levels as assessed by fluorescence intensity (**fig. 1C**). These observations were consistent with a prior report that knocking out PIP4K paralogs elevates PI(4,5)P_2_ levels [25], because PIP4K enzymes can inhibit PIP5Ks independently of their catalytic activity. We therefore reasoned that saturation of endogenous, inhibitory PIP4K molecules by PIP5K over-expression, regardless of catalytic activity of the PIP5K, would free endogenous, active PIP5K enzyme from negative regulation (**fig. 1D**).

**Table 3:** P values from Kruskal-Wallis test with Dunn’s correction for multiple comparisons for Tubbyc^R332H^ (PI(4,5)P_2_) biosensor data presented in Figure 1C. Significant variation was observed among groups by one-way ANOVA (KW statistic = 231.1, P <0.0001). Significant results are highlighted in bold and n values are displayed above.

| Construct | BFP (n=90) | INPP5E | PIP4K2A | PIP4K2A<sup>CD</sup> | PIP4K2B | PIP4K2B<sup>CD</sup> | PIP4K2C |
| --- | --- | --- | --- | --- | --- | --- | --- |
| INPP5E (n=90) | **<0.0001** |  |  |  |  |  |  |
| PIP4K2A (n=90) | **<0.0001** | 0.0032 |  |  |  |  |  |
| PIP4K2A<sup>CD</sup> (n=90) | 0.0658 | **<0.0001** | **0.0159** |  |  |  |  |
| PIP4K2B (n=90) | **<0.0001** | 0.1877 | >0.9999 | **0.0001** |  |  |  |
| PIP4K2B<sup>CD</sup> (n=90) | >0.9999 | **<0.0001** | **<0.0001** | 0.1118 | **<0.0001** |  |  |
| PIP4K2C (n=90) | 0.5904 | **<0.0001** | **0.0008** | >0.9999 | **<0.0001** | 0.8988 |  |
| PIP4K2C<sup>CD</sup> (n=90) | >0.9999 | **<0.0001** | **<0.0001** | **0.0047** | **<0.0001** | >0.9999 | 0.0696 |

To directly test for negative regulation of PIP5K activity by PIP4K in cells, we wanted to assay PI(4,5)P_2_ levels after acute membrane recruitment of normally cytosolic PIP4K paralogs. To this end, we triggered rapid PM recruitment of cytosolic, FKBP-tagged PIP4K by chemically induced dimerization (CID) with a membrane targeted FRB domain, using rapamycin [27]. As shown in **fig. 2A**, all three paralogs of PIP4K induce a steady decline in PM PI(4,5)P_2_ levels within minutes of PM recruitment. Catalytically inactive mutants of all three paralogs produce identical responses (**fig. 2A**).

**Fig. 2.**
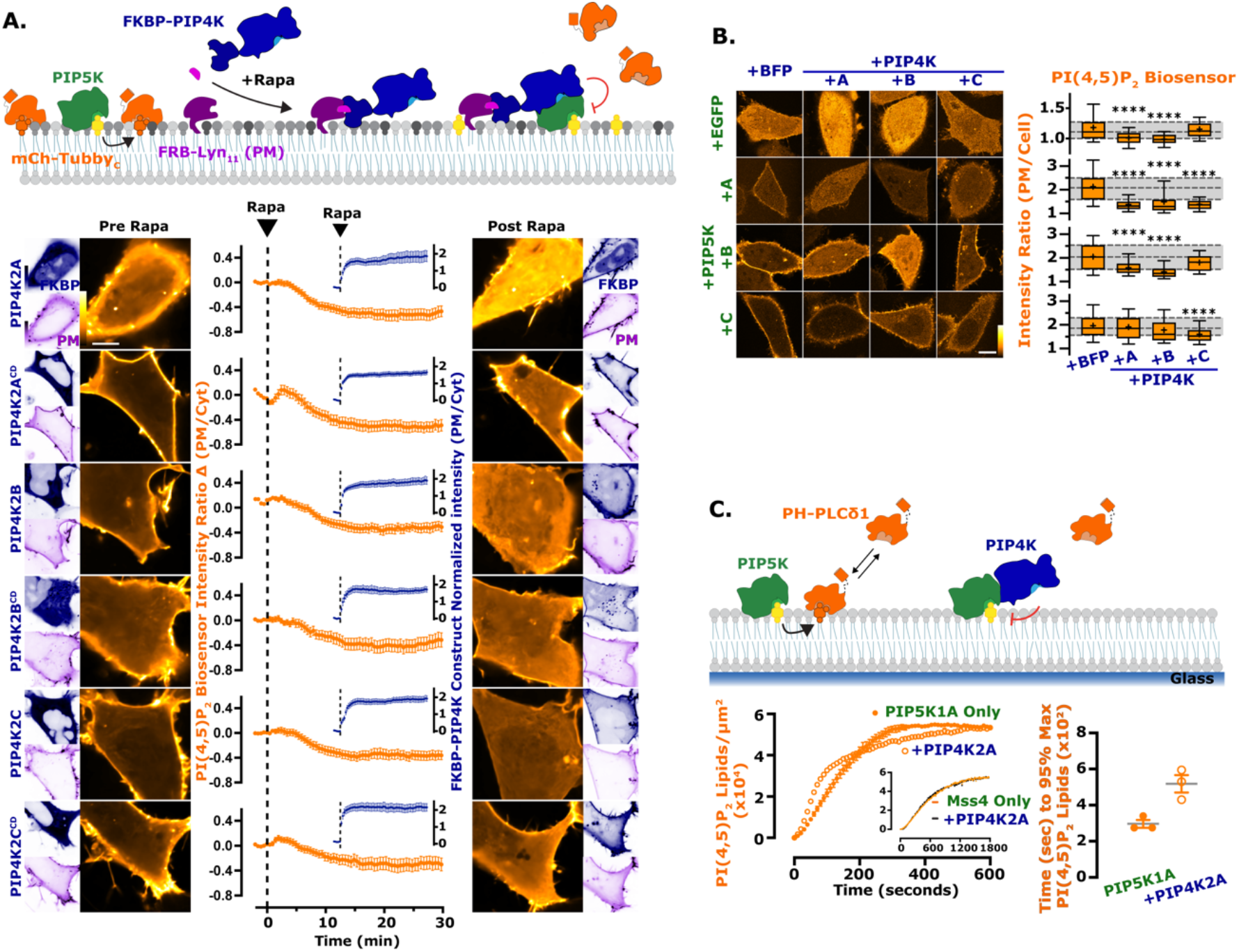
The activity of PIP5K is blunted by PIP4K. (**A**) PIP4K recruitment acutely inhibits PM PI(4,5)P_2_ levels. Cartoon schematics show the chemically induced dimerization (CID) system for FKBP-tagged PIP4K paralogs (A, B, C), which dimerize with the PM-anchored FRB-Lyn11 upon the addition of rapamycin (rapa). HEK293A cells were transfected with FKBP-tagged proteins, the high affinity PI(4,5)P_2_ indicator TubbyC and FRB-Lyn11, scale bar is 10 µm. During time-lapse confocal microscopy, cells were stimulated with 1 µM rapa as indicated. Graphs represent mean change in PI(4,5)P_2_ sensor intensity ratio (PM/Cyt) ± s.e. for 35-60 cells imaged across three independent experiments (orange). Inset graphs show PM recruitment of the FKBP-PIP4K (blue). (**B**) PIP4Ks antagonize PIP5K-mediated PI(4,5)P_2_ increases. HeLa cells expressing PI(4,5)P_2_ indicator TubbyC^R332H^ (orange) were co-transfected with the indicated EGFP- or TagBFP2-tagged constructs. Images show confocal equatorial sections of representative cells, scale bar is 10 µm. Box and whisker plots show the mean fluorescence intensity ratio (PM/Cell) of the PI(4,5)P_2_ sensor from > 90 cells imaged across at least three independent experiments (boxes displaying median and interquartile range, whiskers representing 10-90% of data and “+” represents mean; *P < 0.05; **P < 0.01; ***P < 0.001; ****P < 0.0001). (**C**) PIP4K2A attenuates the kinetics of PI(4,5)P_2_ production driven by PIP5K1A, but not Mss4. Kinetics of PI(4,5)P_2_ production measured on SLBs in the presence of 1 nM PIP5K1A, 20 nM PH-PLCδ1, ± 50 nM PIP4K2A. Inhibition of PIP5K1A activity is delayed until a threshold density of approximately 2% PI(4,5)P_2_ is created to support membrane recruitment of PIP4K2A. Inset shows kinetics of reactions executed in the presence of 50 nM Mss4 , 20 nM PH-PLCδ1, ± 50 nM PIP4K2A. Initial membrane composition: 76% DOPC, 20% DOPS, 4% PI(4)P. Right graphs show the quantification of time required for reactions to reach 95% completion (n = 3 technical replicates).

We also reasoned that co-expression of PIP4K paralogs with PIP5K might attenuate the elevated PI(4,5)P_2_ levels induced by the latter. Broadly speaking, this was true, but with some curious paralog selectivity (**fig. 2B**, statistics reported in **table 4**): PIP4K2A and PIP4K2B both attenuated PI(4,5)P_2_ elevated by PIP5K1A and B, but not (or much less so) PIP5K1C; PIP4K2C, on the other hand, attenuated PIP5K1A and was the only paralog to significantly attenuate PIP5K1C’s effect, yet it did not attenuate PIP5K1B at all.

**Table 4:** P values from Kruskal-Wallis test with Dunn’s correction for multiple comparisons for Tubbyc^R332H^ (PI(4,5)P_2_) biosensor data presented in Figure 2B. Significant variation was observed among groups by one-way ANOVA (FP KW statistic = 74.63, P <0.0001; PIP5K1A KW statistic = 86.53, P <0.0001; PIP5K1B KW statistic = 82.22, P <0.0001; PIP5K1C KW statistic = 23.94, P <0.0001). Significant results are highlighted in bold and n values are displayed.

| EGFP | EGFP (n=90) | PIP4K2A | PIP4K2B |
| --- | --- | --- | --- |
| PIP4K2A (n=90) | **<0.0001** |  |  |
| PIP4K2B (n=90) | **<0.0001** | 0.8805 |  |
| PIP4K2C (n=90) | >0.9999 | **<0.0001** | **<0.0001** |
| PIP5K1A | +BFP | + PIP4K2A | + PIP4K2B |
| + PIP4K2A (n=90) | **<0.0001** |  |  |
| + PIP4K2B (n=90) | **<0.0001** | >0.9999 |  |
| + PIP4K2C (n=90) | **<0.0001** | >0.9999 | >0.9999 |
| PIP5K1B | +BFP | + PIP4K2A | + PIP4K2B |
| + PIP4K2A (n=90) | **<0.0001** |  |  |
| + PIP4K2B (n=90) | **<0.0001** | **0.0027** |  |
| + PIP4K2C (n=90) | 0.7478 | **0.0061** | **<0.0001** |
| PIP5K1C | +BFP | + PIP4K2A | + PIP4K2B |
| + PIP4K2A (n=90) | >0.9999 |  |  |
| + PIP4K2B (n=90) | 0.0933 | 0.3609 |  |
| + PIP4K2C (n=90) | **<0.0001** | **0.0007** | 0.2853 |

To more directly examine inhibition of PIP5K by PIP4K, we tested activity of purified PIP5K1A on PI4P-containing supported lipid bilayers (SLBs). Addition of PIP4K2A exhibited delayed inhibition of PIP5K1A activity (**fig. 2C**): Once PI(4,5)P_2_ reached approximately 28,000 lipids/µm^2^ (∼2 mol %), PIP5K dependent lipid phosphorylation slowed down, which doubled the reaction completion time (**fig. 2C**, right). In contrast, we observed no PIP4K dependent inhibition of Mss4 (**fig. 2C**, inset). These data recapitulate the prior finding that PIP4K only inhibited purified PIP5K in the presence of bilayer-presented substrate[25]. We therefore hypothesized that inhibition of PIP5K by PIP4K requires recruitment of the latter enzyme to the PM by PI(4,5)P_2_ itself.

### PIP4Ks are low affinity sensors of PM PI(4,5)P_2_

To probe the interaction of endogenous PIP4Ks with PM PI(4,5)P_2_, we used a split fluorescent protein genome editing approach [28] to add a NeonGreen2 (NG2) tag to each of the three PIP4K paralogs (**fig. 3A**). Successful integration of the split NG2 tag was evident at the genomic level (**fig. 3B**); a minor shift in protein size was also observed at the protein level after addition of the neonGreen^11^ tag to PIP4K2C (**fig. 3B**). As expected, endogenous PIP4Ks have a mainly cytosolic distribution when viewed in confocal, with a slight enrichment at the cell periphery (**fig. 3C**), which is consistent with results from the OpenCell project [29].

**Fig. 3.**
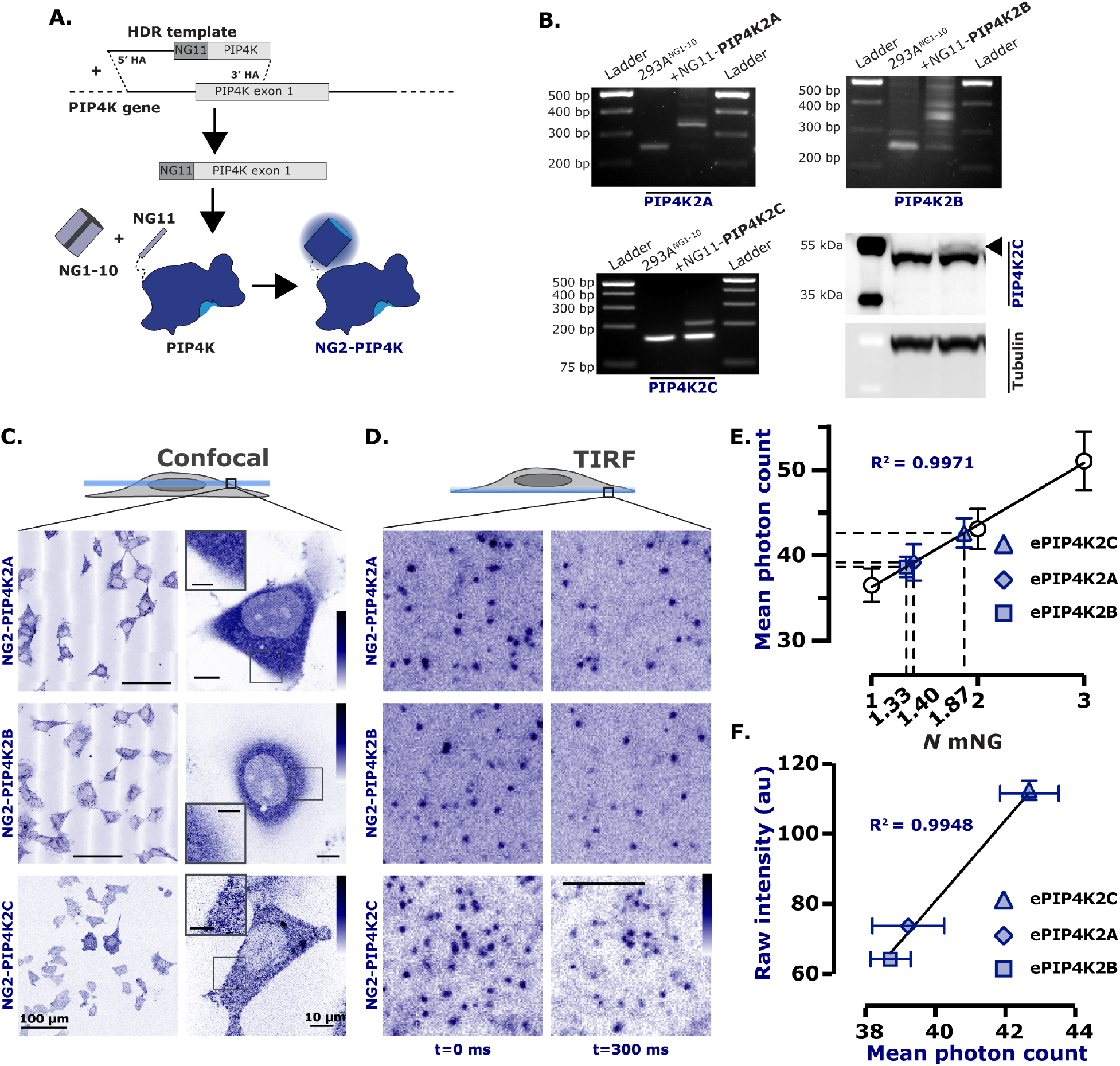
Endogenously tagged PIP4Ks are associated with the PM. (**A**) Endogenous tagging of PIP4K. Brief cartoon schematic showing the mechanism of endogenous tagging employed for PIP4K with NeonGreen2 (NG2). The resulting cell lines were termed NG2-PIP4K. This strategy was used for all three paralogs of PIP4K. (**B**) For PIP4K2A and PIP4K2B paralogs, cells were genotyped with paralog specific forward and reverse primer yielding an edited product of ∼350bp. For PIP4K2C, cells were genotyped with a NG2 specific forward primer and a PIP4K2C paralog specific reverse primer yielding an edited product of ∼200bp. NG2-PIP4K2C cells were also probed with a PIP4K2C specific antibody showing the expected ∼3 kDa shift in weight (arrowhead). (**C**) Confocal based characterization of NG2-PIP4K. Confocal images display the NG2-PIP4K (blue) in cells localized mainly to the cytosol, scale bar is 100 µm or 10 µm, as noted. In the case of NG2-PIP4K2C, slight association of the enzyme to the PM can be seen in the zoomed image, scale bar for zoomed images is 5 µm. (**D**) TIRF based characterization of NG2-PIP4K. When imaged live by TIRF, dynamic, diffraction limited spots are observed on the membrane (compare differential localization at 0 and + 300 ms), scale bar is 2.5 µm. (**E**) Endogenous PIP4Ks exists as heterogeneous populations. All three PIP4K paralogs have an intensity consistent with a mixed population of 1 or 2 mNG molecules when calibrated against single, dimeric or trimeric mNG molecules fused to a PI(4,5)P_2_ binding domain. This correlates to the mean photon count of heterogeneously tagged cell populations (one or two alleles tagged with NG2^11^). Data are grand mean photon counts ± 95% C.I. for data acquired from 22-43 cells. Linear regression and resulting R^2^ against grand mean values is also plotted. (**F**) Endogenous expression levels vary between PIP4K paralogs. The raw fluorescence intensity of NG2-PIP4K2A, NG2-PIP4K2B, or NG2-PIP4K2C in each tagged cell line was measured and plotted against the mean photon counts from E. Plotted pointes show the raw fluorescence intensity of each NG2-PIP4K paralog from 90 cells imaged across three independent experiments (circles display the mean and error bars representing s.e.). There is a positive correlation between the expression level of each paralog and its likelihood to be visualized as a monomeric vs a dimeric protein (Linear regression and resulting R^2^ against mean values is plotted). PIP4K2A and PIP4K2B may exist as either homodimers or more likely heterodimers; whereas PIP4K2C more likely homodimerizes.

Analysis of the ventral PM by total internal reflection fluorescence microscopy (TIRFM) revealed individual, diffraction-limited and uniform intensity puncta that were dynamically associated with the membrane (**fig. 3D**). We compared the intensity of these puncta with a PI(4,5)P_2_ biosensor tagged with single, double or triple mNeonGreen copies expressed at single molecule levels. This revealed that the NG2-PIP4K2C puncta contained an average of 1.87 NG2 molecules, whereas NG2-PIP4K2A puncta contained 1.40 and NG2-PIP4K2B contained 1.33 NG2 molecules. This is consistent with dimeric PIP4K complexes [30] mature copies of the NG2 fluorescent protein and displaying lower fluorescence due to heterodimerization with unlabeled endogenous PIP4Ks (**fig. 3E**). Analysis of the average fluorescence intensity of confocal sections of the edited cells recapitulates the ranked expression order of the PIP4Ks in HEK293 cells by proteomic studies [29,31], with PIP4K2C >> PIP4K2A > PIP4K2B (**fig. 3F**). Satisfyingly, the total intensity of the cells scale linearly with the photon count of single NG2-containing complexes resolved as puncta (**fig. 3F**); this is expected, since PIP4K paralogs exist as a series of randomly associated homo- and hetero dimers of the three paralogs [32]. Therefore, NG2-PIP4K2C dimers are expected to be more frequent since there is more total PIP4K2C expression in HEK293 and thus a higher probability of homodimerization between molecules of this paralog.

Given the dynamic interaction of all three PIP4K paralogs with the PM, we next asked the question: does this interaction depend on PI(4,5)P_2_? On supported lipid bilayers, purified PIP4K2A was released from the membrane upon depletion of PI(4,5)P_2_ by the addition of the 5-OH phosphatase, OCRL (**fig. 4A**), mirroring the kinetics of the PH-PLCδ1 lipid biosensor. To determine if this also holds true with native proteins in living cells, we employed CID to recruit the INPP5E 5-OH phosphatase to rapidly deplete PM PI(4,5)P_2_ [27]. As shown in **figs. 4C-E**, PI(4,5)P_2_ depletion was evident from the rapid loss of a high-affinity biosensor, Tubby_c_ [26]. This depletion was accompanied by loss of PM localized molecules of all three NG2-PIP4K paralogs, PIP4K2A (**fig. 4C**), PIP4K2B (**fig. 4D**) and PIP4K2C (**fig. 4E**) when viewed by TIRFM. Therefore, PI(4,5)P_2_ is necessary to drive PIP4K association with the membrane.

**Fig. 4.**
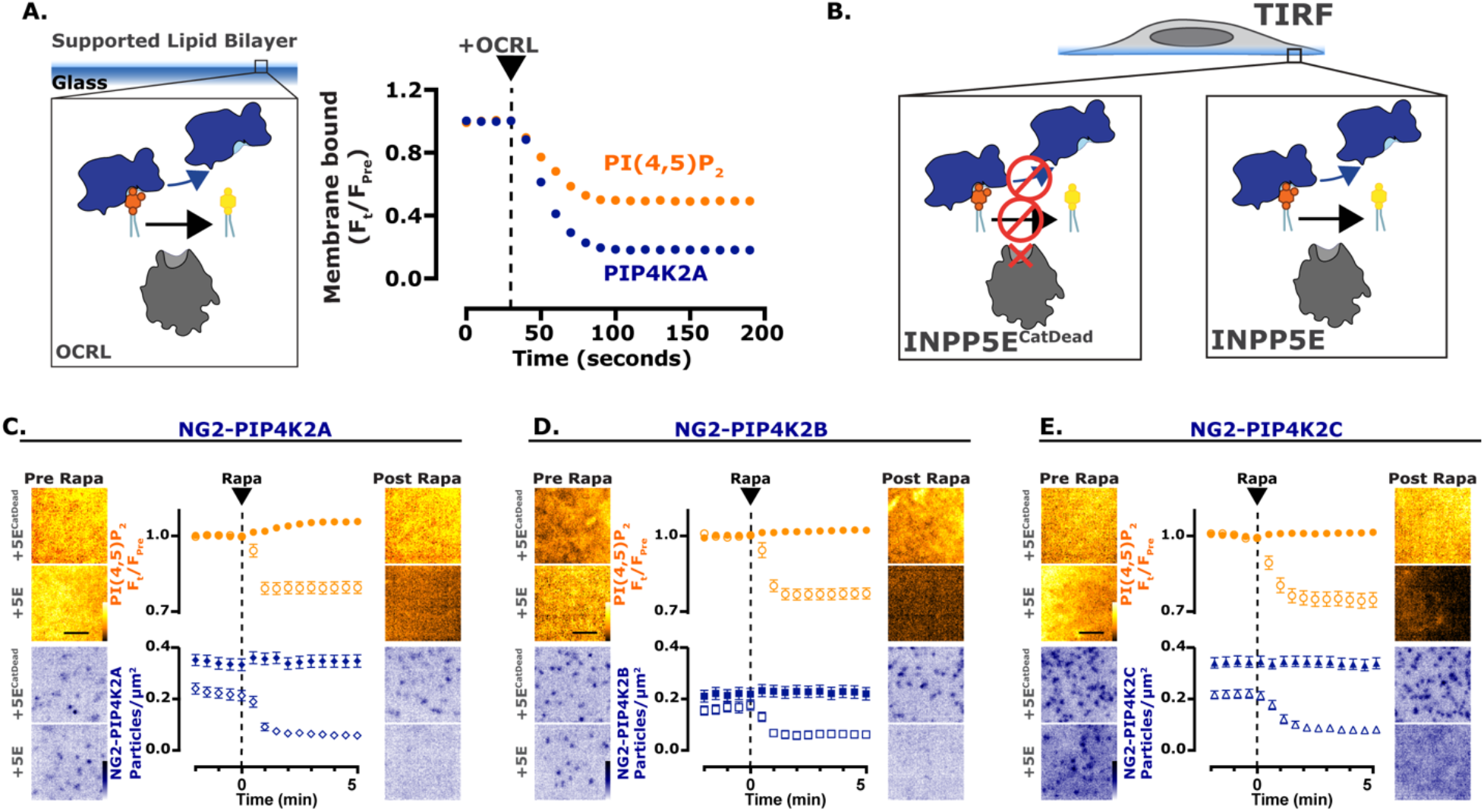
PI(4,5)P_2_ is necessary for the PM localization of PIP4K. (**A**) Depletion of PI(4,5)P_2_ causes PIP4K2A to dissociate from SLBs. Imaging chambers containing 50 nM PIP4K2A and 20 nM PH-PLC81 at equilibrium with SLBs composed of 96% DOPC and 4% PI(4,5)P_2_ were visualized by TIRF microscopy. At 30 seconds, 100 nM OCRL was added to catalyze the dephosphorylation of PI(4,5)P_2_ and membrane dissociation of PIP4K2A and PH-PLC81. (**B**) Depletion of PI(4,5)P_2_ causes NG2-PIP4K2C to dissociate from the membrane. Cartoons show the CID system, in TIRF, for FKBP-tagged INPP5E (catalytically active or dead) dimerizing with the PM-anchored Lyn11-FRB. (**C**) Depletion of PI(4,5)P_2_ causes NG2-PIP4K2A to dissociate from the membrane. NG2-PIP4K2A (blue) cells were transfected with FKBP-tagged proteins, the high affinity PI(4,5)P_2_ indicator TubbyC (orange) and Lyn11-FRB, scale bar is 2.5 µm. During time-lapse TIRF microscopy, cells were stimulated with 1 µM rapa, as indicated. TubbyC traces represent mean change in fluorescence intensity (Ft/Fpre) ± s.e. The NG2-PIP4K2A traces represent the mean change in puncta per µm^2^ ± s.e. of 29-32 cells that were imaged across three independent experiments. (**D**) Depletion of PI(4,5)P_2_ causes NG2-PIP4K2B to dissociate from the membrane. As in C, NG2-PIP4K2B (blue) cells were transfected with FKBP-tagged proteins, TubbyC (orange) and Lyn11-FRB, scale bar is 2.5 µm; cells were stimulated with 1µM rapa, as indicated. TubbyC traces represent mean change in fluorescence intensity (Ft/Fpre) ± s.e. The NG2-PIP4K2B traces represent the mean change in puncta per µm^2^ ± s.e. of > 32 cells that were imaged across three independent experiments. (**E**) Depletion of PI(4,5)P_2_ causes NG2-PIP4K2C to dissociate from the membrane. As in C, NG2-PIP4K2C (blue) cells were transfected with FKBP-tagged proteins, TubbyC (orange) and Lyn11-FRB, scale bar is 2.5 µm; cells were stimulated with 1µM rapa, as indicated. TubbyC traces represent mean change in fluorescence intensity (Ft/Fpre) ± s.e. The NG2-PIP4K2C traces represent the mean change in puncta per µm^2^ ± s.e. of > 38 cells that were imaged across three independent experiments.

Despite this clear PI(4,5)P_2_ binding, a relatively small fraction of PIP4K2C is present on the PM at steady state (see confocal images in **fig. 3C**). Given that there are many orders of magnitude more PI(4,5)P_2_ molecules in the PM than PIP4K in the cell [33], these observations suggest that PIP4Ks bind the lipid with low affinity. Indeed, PIP4K2A binding to supported lipid bilayers was barely evident at 1% PI(4,5)P_2_, but detectable at 2% and rose sharply at 3 and 4% (**fig, 5A**). This is suggestive of a highly co-operative binding mode, as might be expected from a dimeric protein. Notably, binding was not saturated at these low lipid mole fractions, which are thought to be physiological [33]. We therefore reasoned that elevating PM PI(4,5)P_2_ levels may actually increase endogenous PIP4K association with the PM. To this end, we employed over-expression of Mss4, since this enzyme does not bind to PIP4Ks (**fig. 2C**) and enhances PI(4,5)P_2_ in a manner that depends on catalytic activity (**fig. 1A**). Over-expression of Mss4 indeed enhanced membrane binding of all three PIP4K paralogs in a manner dependent on catalytic activity (**fig. 5B**, statistics reported in **table 5**). On PI4P-containing supported lipid bilayers, the addition of active Mss4 induced PIP4K2A binding to the lipid bilayer, again with evidence of co-operativity and a threshold of around 2% PI(4,5)P_2_ (**fig. 5C**).

**Fig. 5.**
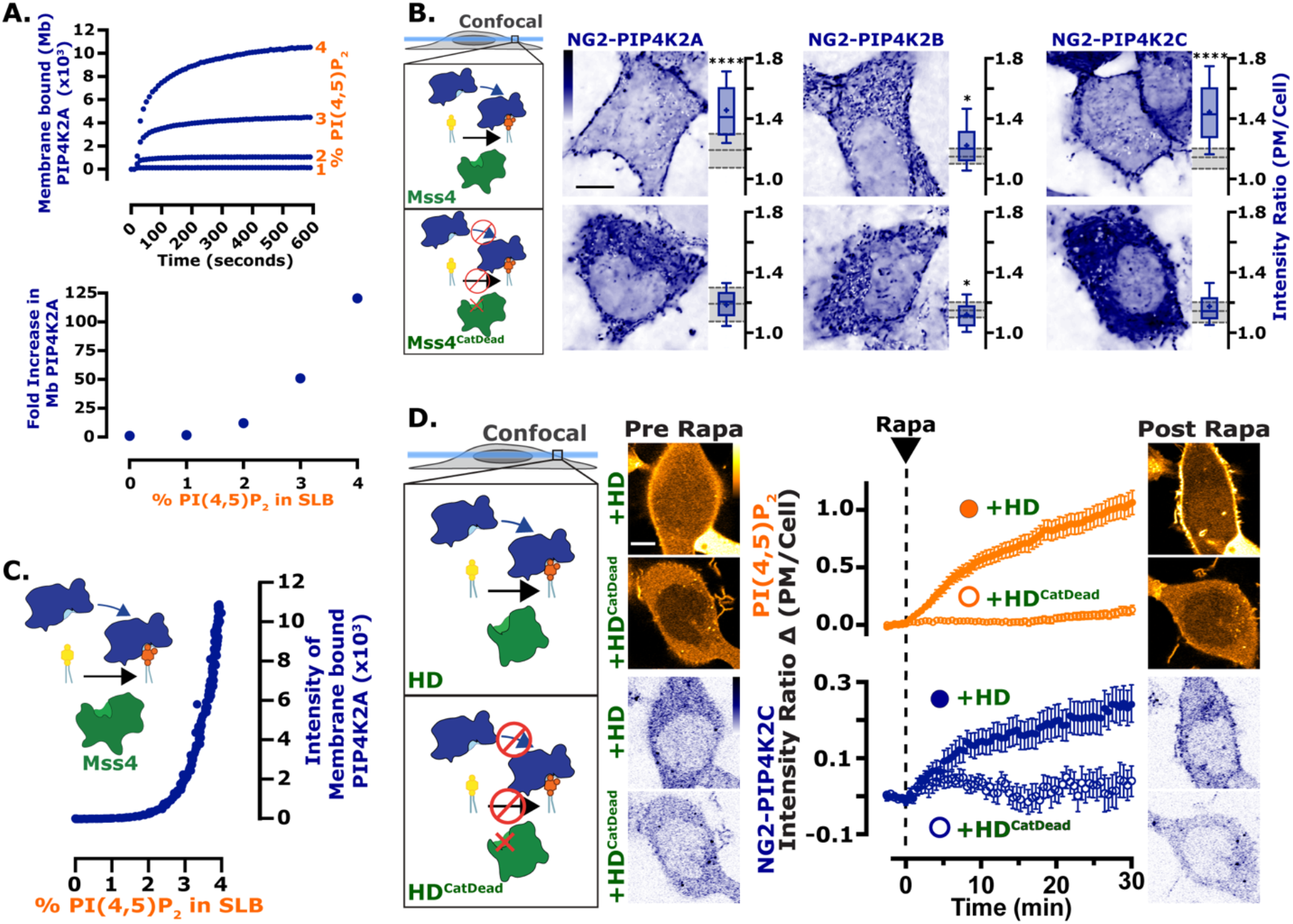
PIP4K is a low affinity PI(4,5)P_2_ sensing protein. (**A**) Purified PIP4K2A localizes to PI(4,5)P_2_ in a concentration dependent manner. Membrane absorption and equilibration kinetics of 50 nM Alexa488-PIP4K2A measured by TIRF microscopy on SLBs containing 1–4% PI(4,5)P_2_. PIP4K2A membrane binding exhibited non-linearity with respect to the PI(4,5)P_2_ lipid density. Quantification of the fold increase in membrane bound PIP4K2A relative to the equilibrium fluorescence intensity of PIP4K2A on a membrane containing 0% PI(4,5)P_2_ is shown in the lower panel. (**B**) Chronic enrichment of PI(4,5)P_2_ causes NG2-PIP4K paralogs to associate with the membrane. Cartoons show the expression of catalytically active or dead Mss4. Images show equatorial confocal sections of representative NG2-PIP4K cells transfected with Mss4, scale bar is 10 µm. Box and whisker plots show the mean fluorescence intensity ratio (PM/Cell) of the indictated NG2-PIP4K paralog from 90 cells images across at least three independent experiments (boxes display median and interquartile range, whiskers represent 10-90% of data and “+” represents mean; *P < 0.05; **P < 0.01; ***P < 0.001; ****P < 0.0001). (**C**) Enrichment of PI(4,5)P_2_ causes dynamic membrane recruitment of purified PIP4K2A. In SLBs, membrane recruitment of 50 nM PIP4K2A monitored during Mss4 catalyzed phosphorylation of PI(4)P. Membranes containing 4% PI(4)P were converted to PI(4,5)P_2_ using 10 nM Mss4. (**D**) Acute enrichment of PI(4,5)P_2_ causes PIP4K2C to increase association with the membrane. Cartoons show the CID system, in confocal, for the interaction of catalytically active or dead FKBP-tagged homo-dimeric PIP5K1C kinase with the PM-anchored Lyn11-FRB. NG2-PIP4K2C (blue) cells were transfected with FKBP-tagged proteins, the low affinity PI(4,5)P_2_ indicator TubbyC^R332H^ (orange) and Lyn11-FRB, scale bar is 10 µm. During time-lapse confocal microscopy, cells were stimulated with 1µM rapa, as indicated. Traces represent mean change in fluorescence intensity (change in PM/Cell ratio from pre-stimulation levels) ± s.e. of 48-52 cells imaged across at least three independent experiments.

**Table 5:** P values from Kruskal-Wallis test with Dunn’s correction for multiple comparisons for indicated NG2-PIP4K paralog data presented in Figure 5B. Significant variation was observed among groups by one-way ANOVA (PIP4K2A KW statistic = 95.78, P <0.0001; PIP4K2B KW statistic = 25.09, P <0.0001; PIP4K2C KW statistic = 96.96, P <0.0001). Significant results are highlighted in bold and n values are displayed.

| NG2-PIP4K2A |  |  | NG2-PIP4K2B |  |  | NG2-PIP4K2C |  |  |
| --- | --- | --- | --- | --- | --- | --- | --- | --- |
|  | BFP (n=90) | Mss4 |  | BFP (n=89) | Mss4 |  | BFP (n=90) | Mss4 |
| Mss4 (n=90) | **<0.0001** |  | Mss4 (n=90) | **0.0313** |  | Mss4 (n=88) | **<0.0001** |  |
| Mss4<sup>CD</sup> (n=90) | >0.9999 | **<0.0001** | Mss4<sup>CD</sup> (n=90) | **0.0448** | **<0.0001** | Mss4<sup>CD</sup> (n=90) | >0.9999 | **<0.0001** |

We next tested for rapid binding to acutely increasing PI(4,5)P_2_ levels in living cells, using CID of a homodimeric mutant PIP5K domain (PIP5K-HD), which can only dimerize with itself and not endogenous PIP5K paralogs [34]. This domain also lacks two basic residues that are crucial for membrane binding [35], and only elevates PM PI(4,5)P_2_ when it retains catalytic activity (**fig. 5D**), unlike the full-length protein (**fig. 1A**). Recruitment of the active mutant PIP5K domain acutely elevated NG2-PIP4K2C membrane association with identical kinetics to the Tubby_c_^R332H^ PI(4,5)P_2_ reporter, whereas the catalytically inactive mutant was without effect (**fig. 5D**).

These data indicate that PIP4K2C binds PM PI(4,5)P_2_ with relatively low affinity. As an additional test of this in live cells, we assessed the kinetics of PM binding during PI(4,5)P_2_ re-synthesis after strong PLC activation. Stimulation of over-expressed PLC-coupled muscarinic M3 receptors induced rapid depletion of both NG2-PIP4K2C and PI(4,5)P_2_ (measured with Tubby_c_, **fig. 6A**). Subsequent induction of PI(4,5)P_2_ re-synthesis with the muscarinic antagonist atropine revealed much slower rebinding of NG2-PIP4K2C to the PM compared to the Tubby_c_ PI(4,5)P_2_ biosensor; PIP4K2C takes more than twice as long (**fig. 6B**).

**Fig. 6.**
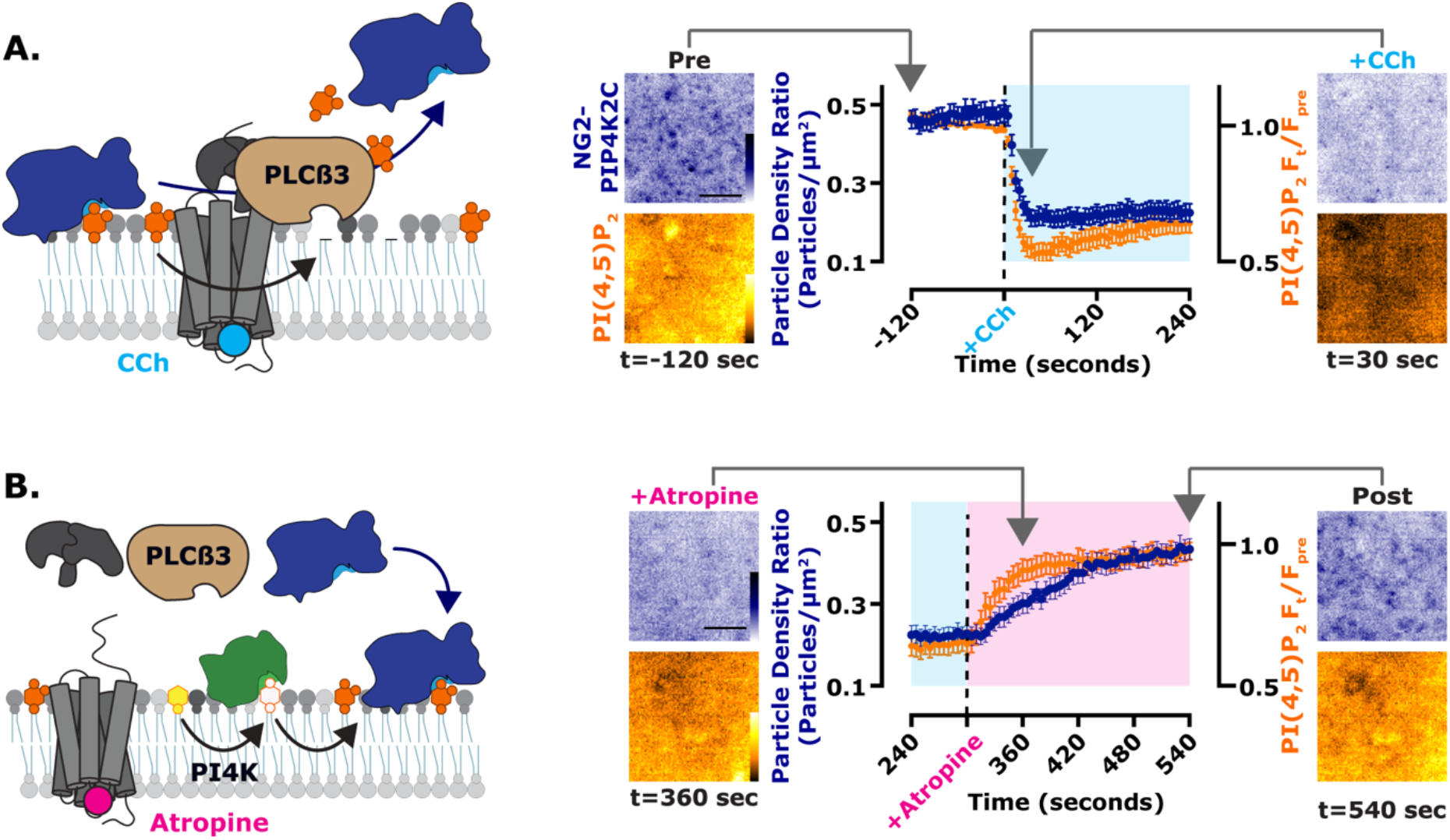
PIP4K binds to the PM at elevated PI(4,5)P_2_ levels. (**A**) PM localization of PIP4K2C follows depletion of PI(4,5)P_2_. Cartoons show PLCβ3 mediated loss of PI(4,5)P_2_ and NG2-PIP4K2C. NG2-PIP4K2C (blue) cells were transfected with the muscarinic acetylcholine receptor M3 and the high affinity PI(4,5)P_2_ indicator TubbyC (orange), scale bar is 2.5 µm. During time-lapse TIRF microscopy, cells were stimulated with 100 µM of the M3 agonist, carbachol (CCh), as indicated. TubbyC traces represent mean change in fluorescence intensity (Ft/Fpre) ± s.e. The NG2-PIP4K traces represent the mean change in puncta per µm^2^ ± s.e. 40 cells were imaged across at least three independent experiments. (**B**) PM localization of PIP4K2C follows resynthesis of PI(4,5)P_2_. Cartoons show the reappearance of PI(4,5)P_2_ and NG2-PIP4K2C after adding the M3 receptor antagonist, atropine. The data are from the later phase of the experiment depicted in **A**. During time-lapse TIRF microscopy, cells were stimulated with 5 µM atropine, as indicated.

Collectively, these data demonstrate that PIP4Ks are low-affinity PI(4,5)P_2_ effectors, poised to sense both decreases and crucially, elevations in PI(4,5)P_2_ levels in the PM. Combined with the previously identified inhibition of PIP5K activity by PIP4K ([25] and **fig. 2**), this suggests a mechanism where PIP4K can act as both receptor and control center for PI(4,5)P_2_ homeostasis, with PIP5K as the effector: when PI(4,5)P_2_ levels rise due to PIP5K activity, PIP4K is recruited to the PM, where it can directly bind and inhibit PIP5K. However, such a mechanism suggests a direct interaction of PIP5K and PIP4K. It is to this question that we turn our attention next.

### PIP4K directly interacts with PIP5K

Previous evidence in the literature points to direct interactions between PIP5Ks and PIP4Ks. Over-expressed PIP4K2A is able to co-immunoprecipitate different PIP5K paralogs [36], and epitope-tagged PIP5K1A was able to pull-down PIP4K2A when expressed at close to endogenous levels [25]. When co-expressing EGFP-tagged PIP5Ks and TagBFP2-tagged PIP4K2s, we found that PIP5K paralogs’ PM binding is largely unaffected by PIP4K over-expression (**fig. 7A**, upper panel and **table 6**), whereas all three paralogs of PIP4K are strongly recruited to the PM by co-expression of any PIP5K (**fig. 7A**, lower panel and **table 7**), as previously observed for PIP4K2A [36].

**Fig. 7.**
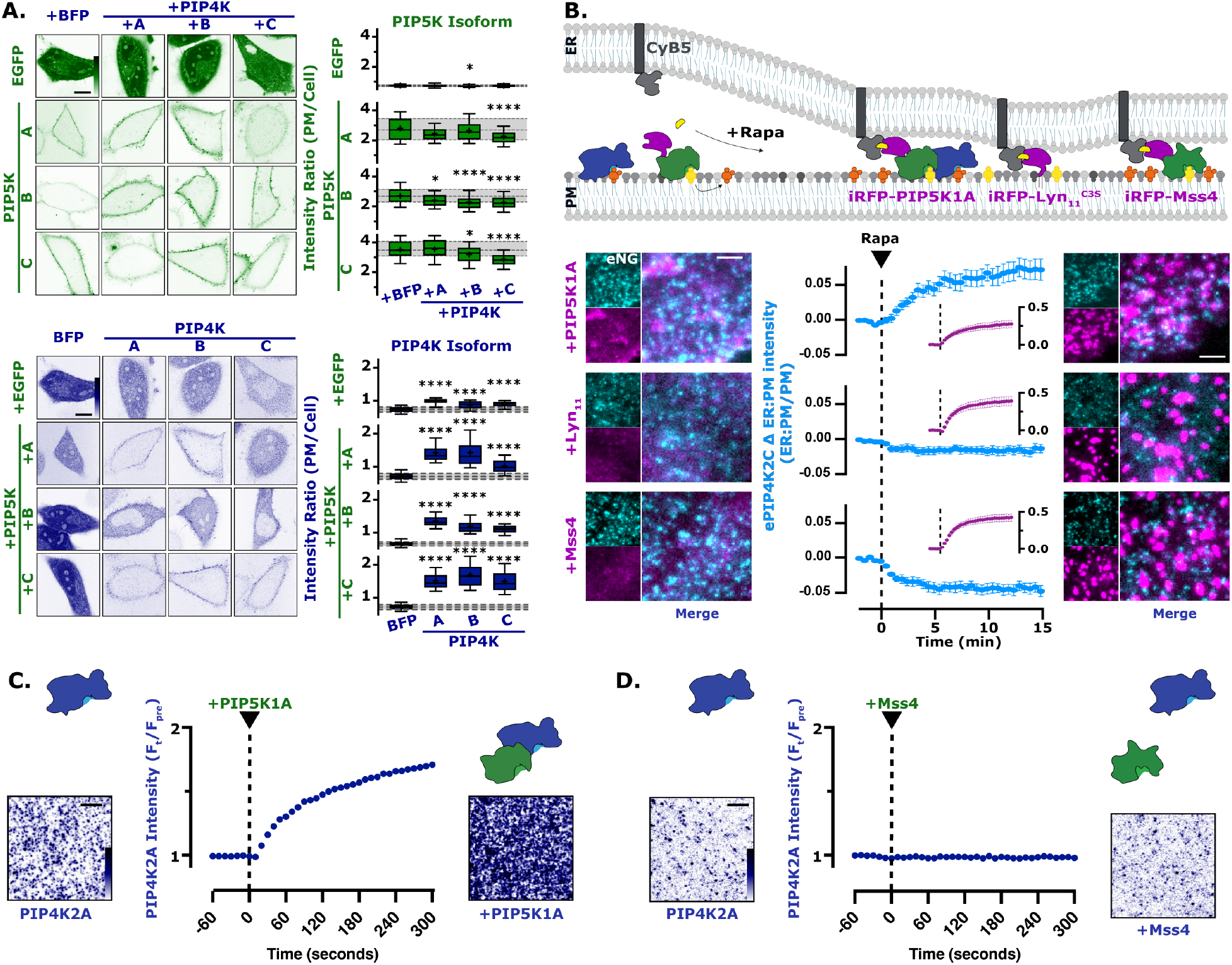
PIP4K directly interacts with mammalian PIP5K. (**A**) PIP5K expression increases PIP4K PM localization. The same experimental data set from Figure 1E is used here. HeLa cells expressing PIP5K (green) or PIP4K (blue) were co-transfected with the indicated EGFP-or TagBFP2-tagged paralog constructs. Images show equatorial sections in confocal of representative cells, scale bar is 10 µm. For box and whisker plots, boxes display median and interquartile range and whiskers representing 10-90% of the data and “+” represents the mean of 90 cells imaged across at least three independent experiments; *P < 0.05; **P < 0.01; ***P < 0.001; ****P < 0.0001. (**B**) PIP4K2C interacts with PIP5K1A. Cartoon schematic show the CID system for the generation of ER-PM contact sites between ER-anchored FKBP-CyB5 and PM-anchored FRB-tagged constructs. NG2-PIP4K2C (cyan) cells were transfected with FKBP-CyB5, mCherry-MAPPER and the indicated FRB-tagged construct (magenta). During time-lapse TIRF microscopy, cells were stimulated with 1µM rapa. TIRF images are representative and color-coded to represent fluorescence intensity, as indicated, scale bar is 2.5 µm. NG2-PIP4K2C traces represent mean fluorescence intensities (ER:PM/PM) ± s.e. of 32-39 cells imaged across a minimum of three independent experiments. (**C**) Dynamic PIP5K1A dependent membrane recruitment of PIP4K2A to SLBs. TIRF microscopy images show the membrane localization of PIP4K2A in the absence and presence of PIP5K1A. In the absence of PIP5K, 50 nM PIP4K2A displays a low level of membrane recruitment. The addition of 10 nM PIP5K1A, stimulates an immediate and steady increase in PIP4K2A membrane localization. Membrane composition: 2% PI(4,5)P_2_, 98% DOPC. TIRF images are representative and color-coded to represent fluorescence intensity, as indicated, scale bar is 5 µm. (**D**) Membrane binding of PIP4K2A is insensitive to yeast Mss4 membrane localization. TIRF microscopy images show the membrane localization of PIP4K2A in the absence and presence of Mss4. Following membrane equilibration of 50 nM PIP4K2A, 10 nM Mss4 was added to the imaging chamber. No appreciable change in PIP4K2A localization was observed during membrane absorption of Mss4. Membrane composition: 2% PI(4,5)P_2_ and 98% DOPC. TIRF images are representative and color-coded to represent fluorescence intensity, as indicated, scale bar is 5 µm.

**Table 6:** P values from Kruskal-Wallis test with Dunn’s correction for multiple comparisons for PIP5K paralogs (green) data presented in the upper panel of Figure 7A. Significant variation was observed among groups by one-way ANOVA (FP KW statistic = 9.161, P = 0.0272; PIP5K1A KW statistic = 25.48, P <0.0001; PIP5K1B KW statistic = 34.88, P <0.0001; PIP5K1C KW statistic = 55.57, P <0.0001). Significant results are highlighted in bold and n values are displayed.

| EGFP | BFP | PIP4K2A | PIP4K2B |
| --- | --- | --- | --- |
| PIP4K2A (n=90) | >0.9999 |  |  |
| PIP4K2B (n=90) | **0.0341** | 0.1387 |  |
| PIP4K2C (n=90) | >0.9999 | >0.9999 | 0.1533 |
| PIP5K1A | +BFP | + PIP4K2A | + PIP4K2B |
| + PIP4K2A (n=90) | 0.0679 |  |  |
| + PIP4K2B (n=90) | >0.9999 | 0.9888 |  |
| + PIP4K2C (n=90) | **<0.0001** | 0.1445 | **0.0016** |
| PIP5K1B | +BFP | + PIP4K2A | + PIP4K2B |
| + PIP4K2A (n=90) | **0.0119** |  |  |
| + PIP4K2B (n=90) | **<0.0001** | 0.3703 |  |
| + PIP4K2C (n=90) | **<0.0001** | 0.1917 | >0.9999 |
| PIP5K1C | +BFP | + PIP4K2A | + PIP4K2B |
| + PIP4K2A (n=90) | >0.9999 |  |  |
| + PIP4K2B (n=90) | **0.0237** | **0.0167** |  |
| + PIP4K2C (n=90) | **<0.0001** | **<0.0001** | **0.0032** |

**Table 7:** P values from Kruskal-Wallis test with Dunn’s correction for multiple comparisons for PIP4K paralogs (blue) biosensor data presented in the lower panel of Figure 7A. Significant variation was observed among groups by one-way ANOVA (FP KW statistic = 136.3, P <0.0001; PIP5K1A KW statistic 222.3, P <0.0001; PIP5K1B KW statistic = 236.6, P <0.0001; PIP5K1C KW statistic = 209.4, P <0.0001). Significant results are highlighted in bold and n values are displayed.

| EGFP | BFP | PIP4K2A | PIP4K2B |
| --- | --- | --- | --- |
| PIP4K2A (n=90) | **<0.0001** |  |  |
| PIP4K2B (n=90) | **<0.0001** | **<0.0001** |  |
| PIP4K2C (n=90) | **<0.0001** | **<0.0001** | >0.9999 |
| PIP5K1A | +BFP | + PIP4K2A | + PIP4K2B |
| + PIP4K2A (n=90) | **<0.0001** |  |  |
| + PIP4K2B (n=90) | **<0.0001** | >0.9999 |  |
| + PIP4K2C (n=90) | **<0.0001** | **<0.0001** | **<0.0001** |
| PIP5K1B | +BFP | + PIP4K2A | + PIP4K2B |
| + PIP4K2A (n=90) | **<0.0001** |  |  |
| + PIP4K2B (n=90) | **<0.0001** | **0.0001** |  |
| + PIP4K2C (n=90) | **<0.0001** | **<0.0001** | 0.5067 |
| PIP5K1C | +BFP | + PIP4K2A | + PIP4K2B |
| + PIP4K2A (n=90) | **<0.0001** |  |  |
| + PIP4K2B (n=90) | **<0.0001** | 0.193 |  |
| + PIP4K2C (n=90) | **<0.0001** | >0.9999 | 0.0544 |

While these data are consistent with a direct interaction between PIP4Ks and PIP5Ks, another possibility exists: the PIP5K dependent increase in PI(4,5)P_2_ (**fig. 1A**) enhances PM recruitment of PIP4K (**figs. 4-6**). Prior pull-downs of PIP5K and PIP4K from lysates required cross-linking the proteins, which may have occurred when the enzymes were simply co-localized on the PM rather than directly interacting [25]. We therefore sought to distinguish between a direct PIP5K-PIP4K binding interaction versus PI(4,5)P_2_-induced co-enrichment on the PM. To this end, we devised an experiment whereby a bait protein (either PIP5K or control proteins) could be acutely localized to subdomains of the PM, with the same PI(4,5)P_2_ concentration. This was achieved using CID of baits with an endoplasmic reticulum (ER) tethered protein, causing restricted localization of the bait protein to ER-PM contact sites – a subdomain of the PM (**fig. 7B**). Enrichment of endogenous NG2-PIP4K2C at ER-PM contact sites was only observed when PIP5K1A was the bait; an unrelated peptide (myristoylated and palmitoylated peptide from Lyn kinase, Lyn_11_) or Mss4 did not enrich NG2-PIP4K2C (**fig. 7B**). The use of Mss4 ruled out an effect of enhanced PI(4,5)P_2_ generation at contact sites, since this enzyme increases PI(4,5)P_2_ as potently as PIP5K1A (**fig. 1A**), yet does not cause recruitment of PIP4K2C.

Finally, we also demonstrate that PIP4K2A binding to PI(4,5)P_2_-containing supported lipid bilayers was greatly enhanced by addition of PIP5K to the membranes (**fig. 7C**), but not by Mss4 (**fig. 7D**). Clearly, PIP4K enzymes directly interact with PIP5Ks on PI(4,5)P_2_-containing lipid bilayers. The ability of PIP4K to bind to PIP5K on a PI(4,5)P_2_-containing bilayer also potentially explains the slightly accelerated initial rate of PI(4,5)P_2_ synthesis exhibited by PIP5K1A that we reported in **fig. 2C**, since PIP4K may initially introduce some avidity to the membrane interaction by PIP5K, before PI(4,5)P_2_ reaches a sufficient concentration that PIP4K-mediated inhibition is effective.

### Disruption of PI(4,5)P_2_ has differential effects on signaling

Synthesizing all of these observations, we propose a simple homeostatic feedback loop that maintains PI(4,5)P_2_ levels in the PM (**fig. 8A**): when PI(4,5)P_2_ levels increase, PIP4K is recruited to the PM in sufficient quantities to inhibit PIP5K, halting further PI(4,5)P_2_ synthesis. If PI(4,5)P_2_ levels fall, PIP4K is one of the first PI(4,5)P_2_ binding proteins to be released (due to its low affinity), causing disinhibition of PIP5K and recovery of PI(4,5)P_2_. We next sought to test how perturbations of this homeostat would affect physiological function. We could produce graded changes in resting PI(4,5)P_2_ levels by over-expression of various components of the homeostat: enhanced PIP5K1A expression, either catalytically active or inactive, increases PI(4,5)P_2_; a myristoylated PIP4K2A retains PM localization even at low PI(4,5)P_2_, causing sustained reductions in PI(4,5)P_2_; and a PM-localized PI(4,5)P_2_ 5-OH phosphatase causes near complete ablation of the lipid. These constructs all show the expected changes in PM PI(4,5)P_2_ compared to a control, reported by three different PI(4,5)P_2_ biosensors. Of these, Tubby_c_ showed the largest degree of change in PM localization across all changes in PI(4,5)P_2_ levels (**fig. 8B**). We then used these graded changes in steady-state PM PI(4,5)P_2_ to investigate the concentration requirements for the lipid in signaling.

**Fig. 8.**
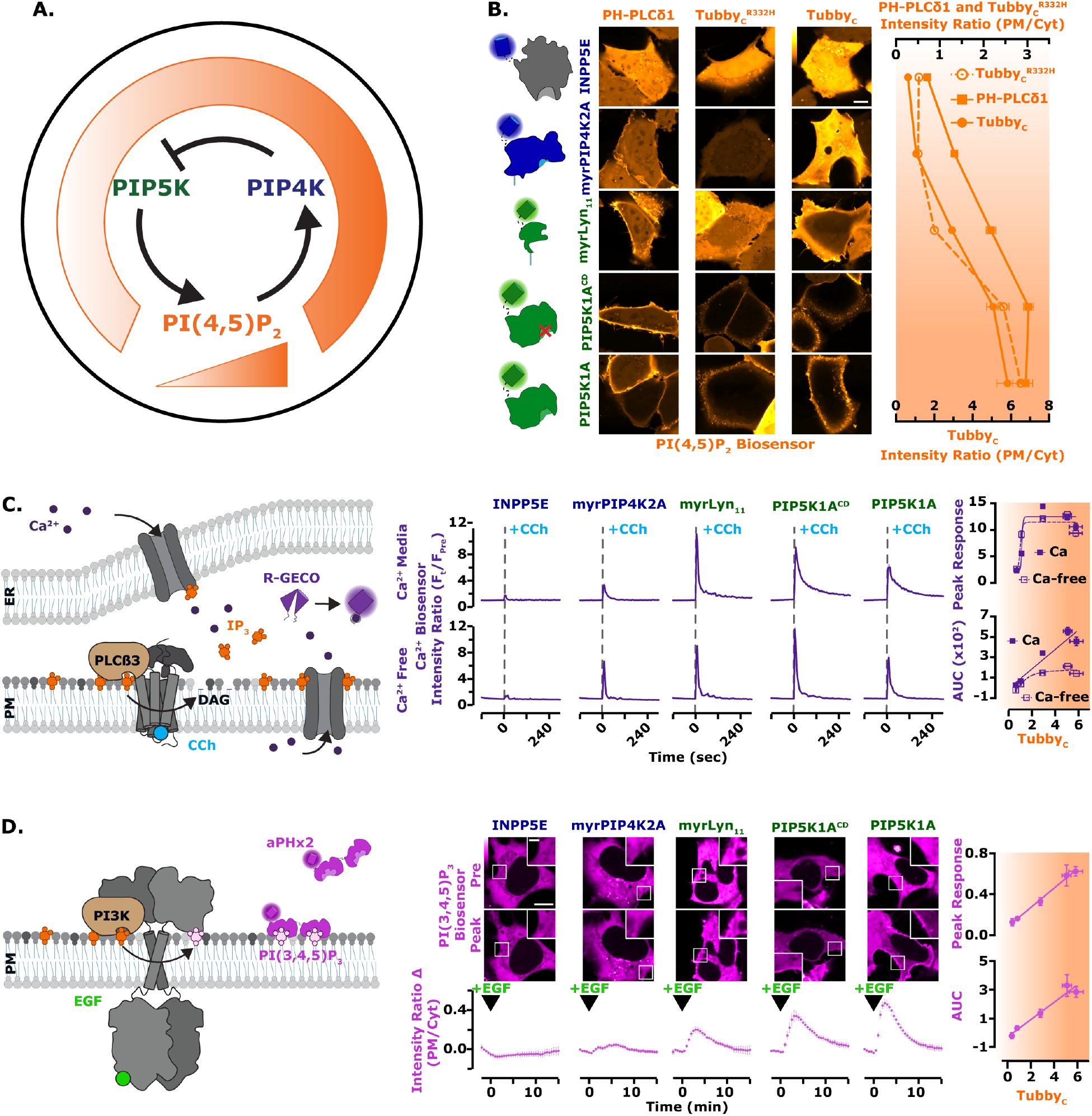
PI3K, but not calcium signaling, are modulated across all concentration ranges of PI(4,5)P_2_. (**A**) Proposed regulation of PIP5K by the low affinity PI(4,5)P_2_ interaction of PIP4K. The working model for negative feedback of PIP5K via PIP4K resembles the thermostat regulation of temperature. When PI(4,5)P_2_ levels are high, PIP4K is recruited and held at the PM, via a direct low affinity interaction with PI(4,5)P_2_. At the PM, PIP4K interacts with and inhibits the catalytic activity of PIP5K, causing reduced PI(4,5)P_2_ synthesis. (**B**) PI(4,5)P_2_ biosensors detect a gradient of lipid levels. HEK293A cells were transfected with the indicated fluorescently tagged PI(4,5)P_2_ modulating proteins (INPP5E, myrPIP4K2A, myrLyn11, PIP5K1A catalytic dead or active) and the indicated PI(4,5)P_2_ biosensor (PH-PLC81, Tubby, or Tubby^R332H^ displayed in orange) for 16-24 hours, scale bar is 10 µm. Mean fluorescence intensity (PM/Cyt) are shown as points with error bars representing s.e. of >120 cells imaged across three independent experiments. (**C**) PLC-mediated Ca^2+^ signals saturate at tonic PI(4,5)P_2_ levels. Cartoon schematics of PLC mediated Ca^2+^ signaling and detection. HEK293A cells were transfected with the indicated fluorescently tagged construct and the calcium sensor R-GECO (purple). During time-lapse confocal microscopy (performed with either compete imaging media containing 1.8 mM Ca^2+^ [Ca^2+^] or calcium free Ringer’s media [Ca^2+^-Free]), cells were stimulated with 100 µM CCh as indicated. Traces represent the peak response of mean change in fluorescence intensity (Ft/FPre normalized to pre-stimulation levels) ± s.e. of >100 cells imaged across a minimum of three independent experiments. The peak response and total area under the curve (AUC) were plotted against the normalized ratio of TubbyC. (**D**) PI3K mediated PI(3,4,5)P_3_ synthesis is linearly dependent on PI(4,5)P_2_ levels. Cartoon schematics show PI3K mediated signaling and detection of PI(3,4,5)P_3_ upon the addition of EGF. HEK293A cells were transfected with the indicated fluorescently tagged construct and the PI(3,4,5)P_3_ biosensor, PH-ARNO^2G-I303Ex2^ (aPHx2) (magenta), scale bar is 10 µm and inset scale bar is 2.5 µm. During time-lapse confocal microscopy, cells were stimulated with 10 ng/mL EGF, as indicated. Traces represent the peak response of mean change in fluorescence intensity (change in PM/Cyt from pre-stimulation levels) ± s.e. of 35 cells imaged across a minimum of three independent experiments. The peak response and AUC were plotted against the normalized ratio of TubbyC.

PI(4,5)P_2_ is the substrate for PLC, the enzyme that cleaves it into second messengers diacylglycerol and inositol (1,4,5)-trisphosphate (IP_3_), triggering calcium release from ER stores (**fig. 8C**). Calcium release was indeed reduced by lower PI(4,5)P_2_ levels, but appeared to be maximal at tonic PI(4,5)P_2_ levels; it was unaffected by increased PM PI(4,5)P_2_. This was true for both peak calcium release and total release from stores (assessed by measuring activity in calcium-free medium, **fig. 8C**). Influx of extracellular calcium was increased by elevated PI(4,5)P_2_ levels (**fig. 8C**), consistent with a prior report that store-operated calcium entry is enhanced by increased PIP5K activity [13]. However, IP_3_-triggered calcium release appears saturated at resting PI(4,5)P_2_. This strongly contrasts with the effects on another PI(4,5)P_2_ signaling pathway, class I phosphoinositide 3-OH kinase (PI3K). Epidermal growth factor (EGF) receptor stimulation activates PI3K, which converts a small fraction of PI(4,5)P_2_ to PIP_3_ (**fig. 8D**). Using a sensitive PIP_3_ biosensor, we observed PIP_3_ production changing proportionately with PI(4,5)P_2_, never reaching a saturated level (**fig. 8D**). PI3K activation therefore, unlike PLC, is sensitive to upregulation by alterations in PI(4,5)P_2_ homeostasis that enhance steady-state levels of the lipid, e.g. by enhanced PIP5K1A expression.

## Discussion

The work presented herein reveals a remarkably simple homeostatic mechanism for PM PI(4,5)P_2_ levels (**fig. 8A**). Here, the PIP4K family of enzymes serve as both receptor and control center, detecting PI(4,5)P_2_ and controlling the activity of the effector, PIP5K. This mechanism is also complementary to a previously identified homeostatic feedback, whereby PI4P catabolism is inactivated in cells until sufficient PI(4,5)P_2_ has been generated [37]. By these mechanisms, cells can ensure adequate PI(4,5)P_2_ is generated to support the cytoskeletal assembly, small solute transport, ion flux, membrane traffic and cell signaling processes controlled by PI(4,5)P_2_. PIP4K’s low affinity and highly co-operative binding to PI(4,5)P_2_ makes it an excellent sensor for tonic PI(4,5)P_2_ levels. It is poised to sense PI(4,5)P_2_ generated in excess of the needs of the lipids’ legion effector proteins, ensuring these needs are met but not exceeded. Nevertheless, the relatively low PIP4K copy number of around 2.5 × 10^5^ per cell [29] is a small fraction of the total PI(4,5)P_2_ pool, estimated to be ∼10^7^ [33], ensuring little impact on the capacity of the lipid to interact with its effectors.

Since this paper was initially submitted for publication, another study has reported a similar homeostatic feedback loop in *Drosophila* photoreceptors, utilizing the fly homologue of septin 7 as the receptor and control center [38]. This conclusion is based on the observation that cells with reduced septin 7 levels have enhanced PIP5K activity in lysates, and exhibit more rapid PI(4,5)P_2_ resynthesis after PLC activation. However, changes in septin 7 membrane localization in response to acute alterations in PI(4,5)P_2_ levels, as well as direct interactions between PIP5K and septin 7, have yet to be demonstrated. Nevertheless, septin 7 has distinct properties as a potential homeostatic mediator; as a foundational member of the septin family, it is essential for generating all major types of septin filament [39]. Therefore, a null allele for this subunit is expected to reduce the prevalence of the septin cytoskeleton by half. Given that septin subunits are found in mammalian cells at high copy number, around ∼10^6^ each [29], and the fact that septins bind PI4P and PI(4,5)P_2_ [40,41], it is likely that septin filaments sequester a significant fraction of the PM PI4P and PI(4,5)P_2_ through high-avidity interactions. In addition, membrane-bound septins appear to be effective diffusion barriers to PI(4,5)P_2_ and other lipids [42]. We therefore speculate that septins may play a unique role in systems such as the fly photoreceptor with extremely high levels of PLC-mediated PI(4,5)P_2_ turnover: The septin cytoskeleton can act as a significant buffer for PI4P and PI(4,5)P_2_ in such systems, as well as corralling pools of the lipids for use at the rhabdomeres were the high rate of turnover occurs. This is in contrast to the role played by the PIP4Ks, where PI(4,5)P_2_ levels are held in a narrow range under conditions of more limited turnover, as found in most cells.

That PIP4K has such a crucial function for which catalytic activity is entirely dispensable is surprising. PIP4K catalytic activity varies among paralogs by almost four orders of magnitude [43]; nevertheless, the ability of the enzymes to phosphorylate PI5P is known to be crucial for many of its other physiological functions [44,45]. However, the low affinity PM PI(4,5)P_2_ binding that we describe, and its inhibition of PIP5K described previously [25], explain why PIP4Ks are expressed in cells in excess of PIP5K by as much as 10:1 [29,31]. This fact does not make sense relative to the enzymes’ catalytic activity, given that PIP4Ks’ substrate, PI5P, is outnumbered by PI4P by around 100-fold [46].

Curiously, although phosphatidylinositol phosphate kinases are found throughout eukarya, PIP4Ks are limited to holozoa (animals and closely related unicellular organisms) [47]. Indeed, we found the PIP5K from the fission yeast, *Saccharomyces cerevisiae*, does not interact with human PIP4Ks (**fig. 7**) and cannot modulate PI(4,5)P_2_ levels in human cells without its catalytic activity (**fig. 1**). This begs the question: how do *S. cerevisiae* regulate their own PI(4,5)P_2_ levels? Intriguingly, they seem to have a paralogous homeostatic mechanism: the dual PH domain containing protein Opy1 serves as receptor and control center [48], in an analogous role to PIP4K. Since there is no mammalian homolog of Opy1, this homeostatic mechanism appears to have appeared at least twice through convergent evolution. Combined with hints of a role for septins in maintaining PI(4,5)P_2_ levels [38], the possibility arises that there may yet be more feedback controls of PI(4,5)P_2_ levels to be discovered.

Despite minor differences in the ability of over-expressed PIP5K paralogs to recruit over-expressed PIP4K enzymes (**fig. 7A**), we observed major differences in the ability of PIP4K paralogs to inhibit PI(4,5)P_2_ synthesis when over-expressed alone (**fig. 1C**) or in combination with PIP5K (**fig. 2B**). It is unclear what drives the partially overlapping inhibitory activity, where each PIP5K paralog can be attenuated by 2 or 3 PIP4Ks. This is however reminiscent of the biology of the PIPKs, where there is a high degree of redundancy among them, with few unique physiological functions assigned to specific paralogs [49]. There may be hints of paralog-specific functions in our data; for example, enhanced PI(4,5)P_2_ induced by over-expressed PIP5K1C is only really attenuated by PIP4K2C (**fig. 2B**). This could imply a requirement for PIP4K2C in regulating PI(4,5)P_2_ levels during PLC-mediated signaling, given the unique requirements for PIP5K1C in this process [50,51]. Regardless, a full understanding of paralog selectivity will need to be driven by a detailed structural analysis of the interaction between PIP4Ks and PIP5Ks - which is not immediately apparent from their known crystal structures, especially since PIP4Ks and PIP5Ks employ separate and distinct dimerization interfaces [49].

The apparently linear dependence of PI3K on available PI(4,5)P_2_ that we revealed after modulating PI(4,5)P_2_ homeostasis (**fig. 8**) explains enhanced PI3K signaling reported in PIP4K-null cells [25,52]. Intriguingly, PIP4Ks were reported to inhibit PI3K/Akt signaling two decades ago, but the mechanism was proposed to be through removal of its PI5P substrate, which was thought to somehow enhance accumulation of PI3K lipid products, PIP_3_ and PI(3,4)P_2_ [53]. The key evidence that it was PI5P that caused the PI3K lipid accumulation came from the observation that it could be recapitulated by the *Shigella flexneri* effector protein IpgD, which generates some PI5P from PI(4,5)P_2_; this and the analogous *Salmonella* effector SopB both activate the PI3K/Akt pathway [53–55]. However, it was recently shown that both SopB and IpgD are in fact novel phosphotransferases that directly convert PI(4,5)P_2_ into the PI3K signaling lipid PI(3,4)P_2_, explaining how these enzymes activate Akt [56]. It therefore seems more likely that PI(4,5)P_2_ downregulation is the most likely explanation for PI3K/Akt pathway inhibition by PIP4Ks.

In conclusion, our results reveal a simple yet elegant homeostatic mechanism that controls PM PI(4,5)P_2_ levels (**fig. 8A**). Perturbation of this homeostasis reveals different sensitivities of PLC and PI3K signaling, with the latter showing enhanced activation with elevated PI(4,5)P_2_. This likely explains why the PI3K, and not the PLC pathway, drives the phenotype of PIP4K-null fruit flies [52]. More broadly, such differences in the sensitivity of PI(4,5)P_2_-dependent PM functions to lipid concentration may go a long way in explaining the phenotypic diversity of diseases associated with dysregulated PI(4,5)P_2_ metabolism. For example, they may explain why a selective inhibitor of PI3Kα can correct aberrant kidney function associated with Lowe syndrome models [9]. Indeed, experimental manipulation of PI(4,5)P_2_ homeostasis will now afford the ability to determine which of the panoply of PI(4,5)P_2_-dependent PM functions are dysregulated by pathological alterations – perhaps bringing novel therapeutic targets into view.

## Materials and Methods

### Cell culture and lipofection

HeLa (ATCC CCL-2) and HEK293A (ThermoFisher R705-07) cells were cultured in DMEM (low glucose; Life Technologies 10567022) supplemented with 10% heat-inactivated fetal bovine serum (Life Technologies 10438-034), 100 units/ml penicillin, 100µg/ml streptomycin (Life Technologies 15140122), and 1:1,000 chemically defined lipid supplement (Life Technologies 11905031) at 37°C with a humidified atmosphere with 5% CO_2_. Cells were passaged twice per week diluting 1 in 5 after dissociation in TrpLE (Life Technologies 12604039). 293A cells with endogenous PIP4K2 paralog alleles tagged with split NeonGreen2 (NG2) were generated similarly as described [57] using a protocol we have described [58]. In brief, Platinum Cas9 (Thermo Fisher B25640) was precomplexed with gRNA and electroporated into HEK293^NG2-1-10^ cells in combination with a single-stranded HDR Template (IDT). Sequences are provided in **Table 9**. The HDR template contains 70 bp homology-arms, the NG2-11 sequence, and a flexible linker in frame with the appropriate PIP4K paralog: PIP4K2A and PIP4K2B (CATCATATCGGTAAAGGCCTTTTGCCACTCCTTGAAGTTGAGCTCGG TACCACT TCCTGGACCTTGAAACAAAACTTCCAATCCGCCACC ) and PIP4K2C (ATGACCGAGCTCAACTTCAAGGAGTGGCAAAAGGCCTTTACCGATATGATGGGTGGCGGC). After recovery, FACS (University of Pittsburgh Flow Cytometry Core) was used to sort NG2-positive cells. These NG2-PIP4K2A, PIP4K2B, and PIP4K2C cells were cultured under identical conditions to the HeLa and HEK293A cells.

### Chemicals and reagents

Rapamycin (Thermo Fisher BP2963-1) was dissolved in DMSO at 1 mM and stored as a stock at -20°C, it was used in cells at 1 µM. EGTA (VWR EM-4100) was dissolved in water at 0.5 M and stored at room temperature, it was used in cells at 5 mM. EGF (Corning CB-40052) was dissolved in water at 100 µg/ml and stored as a stock at -20°C, it was used in cells at 10 ng/ml. Carbachol (Thermo Fisher AC10824-0050) was dissolved in water at 50 mM and stored as a stock at -20°C, it was used in cells at 100 µM. Atropine (Thermo Fisher AC226680100) was dissolved in 100% ethanol at 25 mM and stored as a stock at -20°C, it was used in cells at 5 µM.

### Plasmids and cloning

The EGFP (*Aequorea victoria* GFP containing F64L and S65T mutations) [59], mCherry (*Discoma* DsRed monomeric variant)[60], mTagBFP_2_ (*Entacmaea quadricolor* protein eqFP578)[61], iRFP713 (*Rhodopseudomonas palustris* [Rp] bacteriophytochrome BphP2)[62] and iRFP670 (RpBphP6 iRFP702 containing V112I, K174M and I247C mutations) [63] fluorophores were used in the Clontech pEGFP-C1, -C2, and -N1 backbones as described previously [58]. Mutated constructs were generated using site-directed mutagenesis using targeted pairs of DNA oligos which were custom made and supplied by Thermo Fisher. New plasmids used in this study were generated using standard restriction-ligation or by using NEBuilder HiFi DNA Assembly (New England Biolabs E552OS). HsPIP5K1A, HsPIP5K1B, Mss4, and HsPIP4K2C were obtained as human codon optimized synthetic gBlocks (IDT). Otherwise, plasmids were obtained from the sources listed in **Table 8**. All constructs were sequence verified using Sanger DNA sequencing. Plasmids constructed for this study are available through Addgene.

**Table 8.**
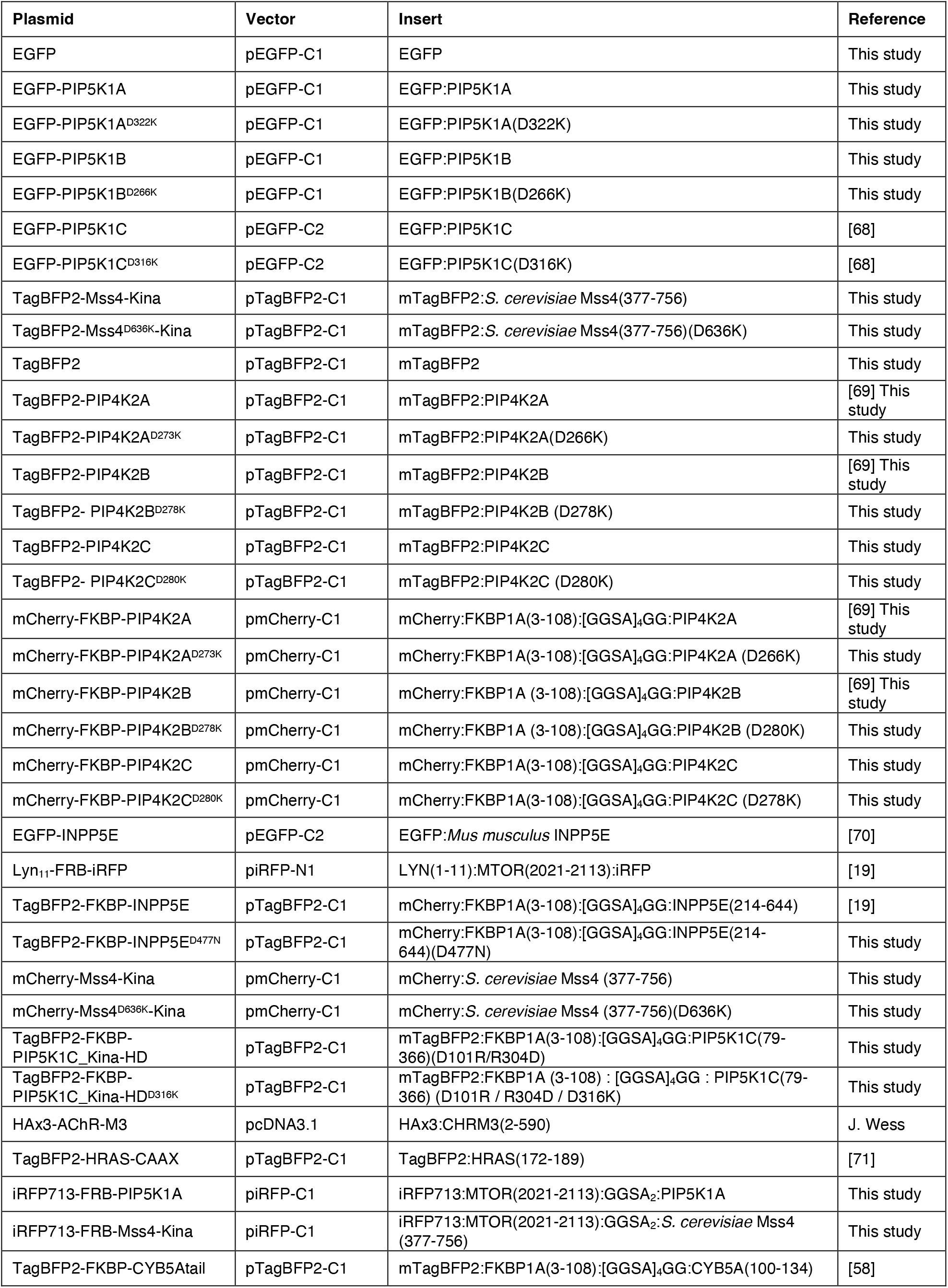
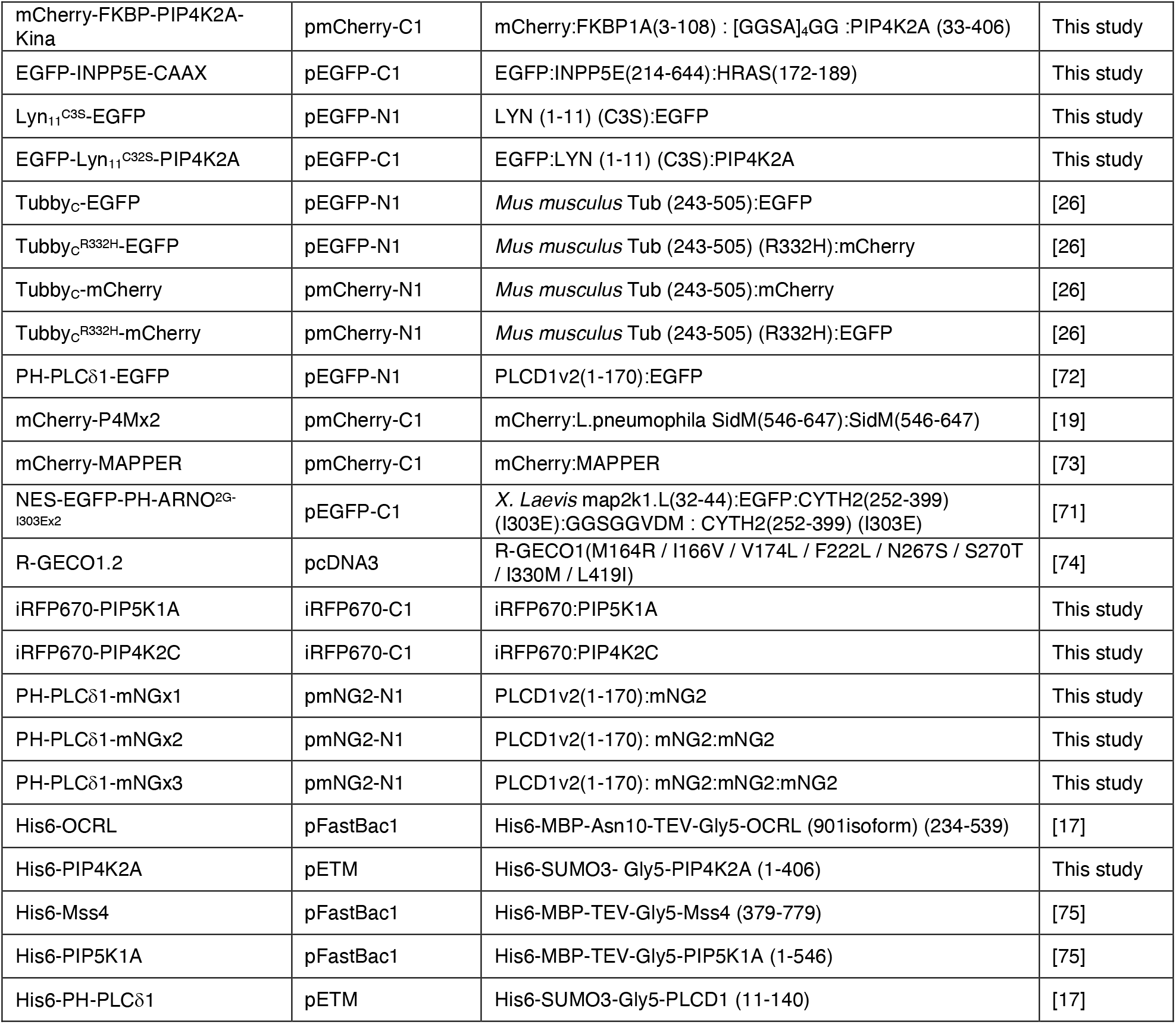
Plasmids used in this study.

**Table 9.**
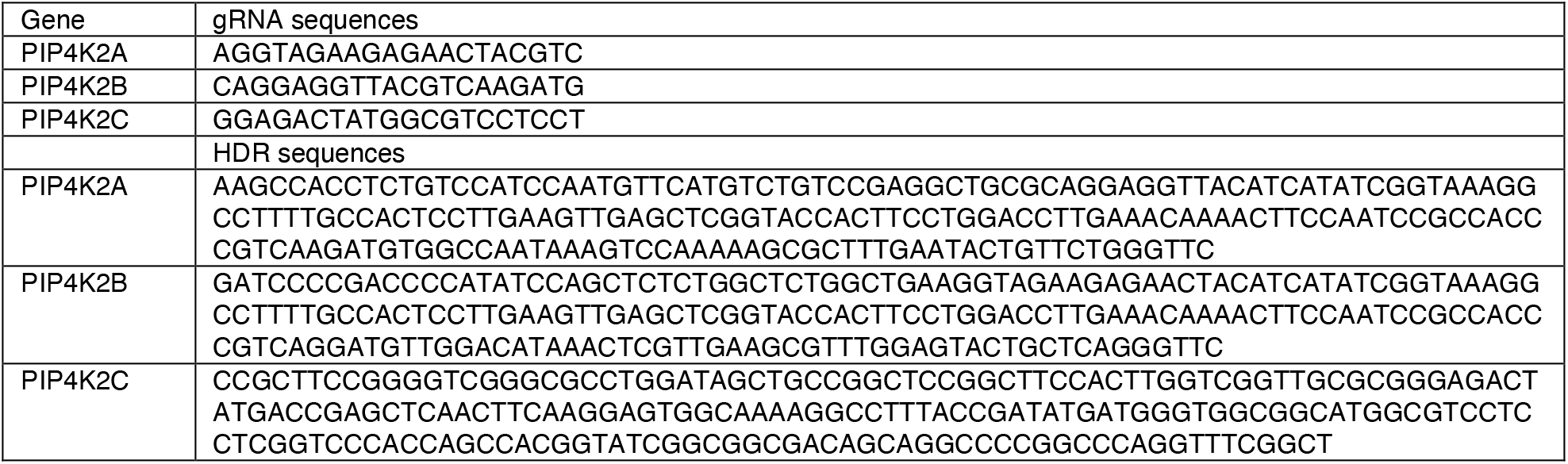
HDR and gRNA sequences for PIP4K2C.

### Purification of PIP5K1A and Mss4

Gene sequences encoding human PIP5K1A and yeast Mss4 kinase domain were cloned into a FastBac1 vector to create the following vectors: His6-MBP-TEV-(Gly)5-PIP5K1A (1-546aa) and His6-MBP-TEV-(Gly)5-Mss4 (379-779aa). BACMIDs and baculovirus were generated as previously described [17]. ES-Sf9 cells were infected with baculovirus using an optimized multiplicity of infection (MOI), typically 2% vol/vol, that was empirically determined from small-scale test expression. Infected cells were typically grown for 48 hours at 27°C in ESF 921 Serum-Free Insect Cell Culture medium (Expression Systems, Cat# 96-001-01) and then harvested by centrifugation. Insect cell pellets were then washed with 1x PBS [pH 7.2] and centrifuged (3500 rpm for 10 minutes). The final cell pellet was combined with an equal volume of buffer containing 1x PBS [pH 7.2], 10% glycerol, and 2x Sigma protease inhibitor cocktail tablet solution before transferring to the -80°C freezer for storage. For purification, frozen cells were thawed in an ambient water bath and then resuspended in buffer containing 50 mM Na_2_HPO_4_ [pH 8.0], 10 mM imidazole, 400 mM NaCl, 5% glycerol, 1 mM PMSF, 5 mM BME, 100 µg/mL DNase, and 1x Sigma protease inhibitor cocktail tablet. Cells were lysed using a glass dounce homogenizer. Lysate was then centrifuged at 35,000 rpm (140,000 × g) for 60 minutes in a Beckman Ti-45 rotor at 4°C. High speed supernatant was combined with 6 mL of Ni-NTA Agarose (Qiagen, Cat# 30230) and stirred in a beaker for 1-2 hour(s) at 4°C. Following batch binding, resin was collected in 50 mL tubes, centrifuged, and washed with buffer containing 50 mM Na_2_HPO_4_ [pH 8.0], 10 mM imidazole, 400 mM NaCl, and 5 mM BME. Ni-NTA resin with His6-MBP-(Asn)10-TEV-(Gly)5-PIP5KA bound was washed in a gravity flow column with 100 mL of 50 mM Na_2_HPO_4_ [pH 8.0], 30 mM imidazole, 400 mM NaCl, 5% glycerol, and 5 mM BME buffer. Protein elution was achieved by washing the resin with buffer containing 50 mM Na_2_HPO_4_ [pH 8.0], 500 mM imidazole, 400 mM NaCl, 5% glycerol, and 5 mM BME. Peak fractions were pooled, combined with 200 µg/mL His6-TEV(S291V) protease, and dialyzed against 4 liters of buffer containing 20 mM Tris [pH 8.0], 200 mM NaCl and 2.5 mM BME for 16-18 hours at 4°C. The next day, dialysate was combined 1:1 volumes with 20 mM Tris [pH 8.0], 1 mM TCEP to reduce the NaCl to a final concentration of 100 mM. Precipitate was removed by centrifugation (3500 rpm for 10 minutes) and a 0.22 µm syringe filtration. Clarified dialysate was bound to a MonoS cation exchange column (GE Healthcare, Cat# 17-5168-01) equilibrated with buffer containing 20 mM Tris [pH 8.0], 100 mM NaCl, and 1 mM TCEP. Proteins were resolved over a 10-100% linear gradient (0.1-1 M NaCl, 45 CV, 45 mL total, 1 mL/min flow rate). (Gly)_x5_-PIP5K1A and (Gly)_x5_-Mss4 eluted from the MonoS in the presence of 375-450 mM NaCl. Peak fractions containing PIP5K1A were pooled, concentrated in a 30 kDa MWCO Vivaspin 6 centrifuge tube (GE Healthcare, Cat# 28-9323-17), and loaded onto a 24 mL Superdex 200 10/300 GL (GE Healthcare, Cat# 17-5174-01) size exclusion column equilibrated in 20 mM Tris [pH 8.0], 200 mM NaCl, 10% glycerol, 1 mM TCEP. Peak fractions were concentrated to 10-50 µM using a 30 kDa MWCO Amicon centrifuge tube (Millipore Sigma) before snap freezing with liquid nitrogen. PIP5K1A and Mss4 were stored in - 80°C as single use aliquots.

### Purification of PIP4K2A

The gene encoding human PIP4K2A was cloned into a pETM derived bacterial expression vector to create the following fusion protein: His6-SUMO3-(Gly)5-PIP4K2A (1-406aa). Recombinant PIP4KA was expressed in BL21 (DE3) Star E. coli (i.e. lack endonuclease for increased mRNA stability). Using 4 liters of Terrific Broth, bacterial cultures were grown at 37°C until OD_600_=0.6. Cultures were then shifted to 18°C for 1 hour to cool down. Protein expression was induced with 50 µM IPTG and bacteria expressed protein for 20 hours at 18°C before being harvested by centrifugation. For purification, cells were lysed into buffer containing 50 mM Na_2_HPO_4_ [pH 8.0], 400 mM NaCl, 0.4 mM BME, 1 mM PMSF (add twice, 15 minutes intervals), DNase, and 1 mg/mL lysozyme using a microtip sonicator. Lysate was centrifuged at 16,000 rpm (35,172 × g) for 60 minutes in a Beckman JA-17 rotor chilled to 4°C. Lysate was circulated over 5 mL HiTrap Chelating column (GE Healthcare, Cat# 17-0409-01) that had been equilibrated with 100 mM CoCl_2_ for 1 hour, washed with MilliQ water, and followed by buffer containing 50 mM Na_2_HPO_4_ [pH 8.0], 400 mM NaCl, 0.4 mM BME. Recombinant PIP4K2A was eluted with a linear gradient of imidazole (0-500 mM, 8 CV, 40 mL total, 2 mL/min flow rate). Peak fractions were pooled, combined with 50 µg/mL of His6-SenP_2_ (SUMO protease), and dialyzed against 4 liters of buffer containing 25 mM Na_2_HPO_4_ [pH 8.0], 400 mM NaCl, and 0.4 mM BME for 16-18 hours at 4°C. Following overnight cleavage of the SUMO3 tag, dialysate containing His6-SUMO3, His6-SenP2, and GGGGG-PIP4K2A was recirculated for at least 1 hour over a 5 mL HiTrap(Co^2+^) chelating column. Flow-through containing GGGGG-PIP4K2A was then concentrated in a 30 kDa MWCO Vivaspin 6 before loading onto a Superdex 200 size exclusion column equilibrated in 20 mM HEPES [pH 7], 200 mM NaCl, 10% glycerol, 1 mM TCEP. In some cases, cation exchange chromatography was used to increase the purity of GGGGG-PIP4K2A before loading on the Superdex 200. In those cases, we equilibrated a MonoS column with 20 mM HEPES [pH 7], 100 mM NaCl, 1 mM TCEP buffer. PIP4K2A (pI = 6.9) bound to the MonoS was resolved over a 10-100% linear gradient (0.1-1 M NaCl, 30 CV, 30 mL total, 1.5 mL/min flow rate). Peak fractions collected from the Superdex 200 were concentrated in a 30 kDa MWCO Amicon centrifuge tube and snap frozen at a final concentration of 20-80 µM using liquid nitrogen.

### Purification of PH-PLCδ1 domain

The coding sequence of human PH-PLCδ1 (11-140aa) was expressed in BL21 (DE3) E. coli as a His6-SUMO3-(Gly)5-PLCδ1 (11-140aa) fusion protein. Bacteria were grown at 37°C in Terrific Broth to an OD_600_ of 0.8. Cultures were shifted to 18°C for 1 hour, induced with 0.1 mM IPTG, and allowed to express protein for 20 hours at 18°C before being harvested. Cells were lysed into 50 mM Na_2_HPO_4_ [pH 8.0], 300 mM NaCl, 0.4 mM BME, 1 mM PMSF, 100 µg/mL DNase using a microfluidizer. Lysate was then centrifuged at 16,000 rpm (35,172 × g) for 60 minutes in a Beckman JA-17 rotor chilled to 4°C. Lysate was circulated over 5 mL HiTrap Chelating column (GE Healthcare, Cat# 17-0409-01) charged with 100 mM CoCl_2_ for 1 hour. Bound protein was then eluted with a linear gradient of imidazole (0-500 mM, 8 CV, 40 mL total, 2 mL/min flow rate). Peak fractions were pooled, combined with SUMO protease (50 µg/mL final concentration), and dialyzed against 4 liters of buffer containing 50 mM Na_2_HPO_4_ [pH 8.0], 300 mM NaCl, and 0.4 mM BME for 16-18 hours at 4°C. Dialysate containing SUMO cleaved protein was recirculated for 1 hour over a 5 mL HiTrap Chelating column. Flow-through containing (Gly)_5_-PLCδ1 (11-140aa) was then concentrated in a 5 kDa MWCO Vivaspin 20 before being loaded on a Superdex 75 size exclusion column equilibrated in 20 mM Tris [pH 8.0], 200 mM NaCl, 10% glycerol, 1 mM TCEP. Peak fractions containing (Gly)_5_-PLCδ1 (11-140aa) were pooled and concentrated to a maximum of 75 µM (1.2 mg/mL) before freezing in liquid nitrogen.

### Purification of OCRL

The coding sequence of human 5-phosphatase OCRL (234-539aa of 901aa isoform) was expressed in BL21 (DE3) E. coli as a His6-MBP-(Asn)10-TEV-(Gly)5-OCRL fusion protein. Bacteria were grown at 37°C in Terrific Broth to an OD600 of 0.8. Cultures were shifted to 18°C for 1 hour, induced with 0.1 mM IPTG, and allowed to express protein for 20 hours at 18°C before being harvested. Cells were lysed into 50 mM Na H_2_PO_4_ [pH 8.0], 300 mM NaCl, 0.4 mM BME, 1 mM PMSF, 100 µg/mL DNase using a microfluidizer. Lysate was then centrifuged at 16,000 rpm (35,172 × g) for 60 minutes in a Beckman JA-17 rotor chilled to 4°C. Lysate was circulated over 5 mL HiTrap Chelating column (GE Healthcare, Cat# 17-040901) charged with 100 mM CoCl2 for 1 hour. Bound protein was eluted with a linear gradient of imidazole (0-500 mM, 8 CV, 40 mL total, 2 mL/min flow rate). Peak fractions were pooled, combined with TEV protease (75 µg/mL final concentration), and dialyzed against 4 liters of buffer containing 50 mM Na H_2_PO_4_ [pH 8.0], 300 mM NaCl, and 0.4 mM BME for 16-18 hours at 4°C. Dialysate containing TEV protease cleaved protein was recirculated for 1 hour over a 5 mL HiTrap Chelating column. Flow-through containing (Gly)5-protein was then concentrated in a 5 kDa MWCO Vivaspin 20 before being loaded on a Superdex 75 (10/300 GL) size exclusion column equilibrated in 20 mM Tris [pH 8.0], 200 mM NaCl, 10% glycerol, 1 mM TCEP. Peak fractions were pooled and concentrated before snap freezing in liquid nitrogen.

### Sortase mediated peptide ligation

PIP4K2A, PIP5K1A, and PH-PLCδ1 were labeled on a N-terminal (Gly)5 motif using sortase mediated peptide ligation[17,64]. Initially, a LPETGG peptide was labeled with either Alexa488, Alexa647, or Cy5 conjugated to an amine reactive N-Hydroxysuccinimide (NHS) (e.g. NHS-Alexa488). Protein labeling was achieved by combining the fluorescently labeled LPETGG peptide with the following reagents: 50 mM Tris [pH 8.0], 150 mM NaCl, 50 µM (Gly)_5_-protein, 500 µM Alexa488-LPETGG, and 10-15 µM His_6_-Sortase. This reaction mixture was incubated at 16-18°C for 16-20 hours, before buffer exchange with a G25 Sephadex column (e.g. PD10) to remove the majority of dye and dye-peptide. The His_6_-Sortase was then captured on Ni-NTA agarose resin (Qiagen) and unbound labeled protein was separated from remaining fluorescent dye and peptide using a Superdex 75 or Superdex 200 size exclusion column (24 mL bed volume).

### Preparation of small unilamellar vesicles

The following lipids were used to generated small unilamellar vesicles (SUVs): 1,2-dioleoyl-sn-glycero-3-phosphocholine (18:1 DOPC, Avanti # 850375C), L-α-phosphatidylinositol-4-phosphate (Brain PI(4)P, Avanti Cat# 840045X), L-α-phosphatidylinositol-4,5-bisphosphate (Brain PI(4,5)P_2_, Avanti # 840046X), and 1,2-dioleoyl-sn-glycero-3-phospho-L-serine (18:1 DOPS, Avanti # 840035C). Lipids were purchased as single use ampules containing between 0.1-5 mg of lipids dissolved in chloroform. Brain PI(4)P and PI(4,5)P_2_ were purchased as 0.25 mg/mL stocks dissolved in chloroform:methanol:water (20:9:1). To make liposomes, 2 µmoles total lipids were combined in a 35 mL glass round bottom flask containing 2 mL of chloroform. Lipids were dried to a thin film using rotary evaporation with the glass round-bottom flask submerged in a 42°C water bath. After evaporating all the chloroform, the round bottom flask was flushed with nitrogen gas for at least 30 minutes. We resuspended the lipid film in 2 mL of PBS [pH 7.2], making a final concentration of 1 mM total lipids. All lipid mixtures expressed as percentages (e.g. 98% DOPC, 2% PI(4)P) are equivalent to molar fractions. For example, a 1 mM lipid mixture containing 98% DOPC and 2% PI(4)P is equivalent to 0.98 mM DOPC and 0.02 mM PI(4)P. To generate 30-50 nm SUVs, 1 mM total lipid mixtures were extruded through a 0.03 µm pore size 19 mm polycarbonate membrane (Avanti #610002) with filter supports (Avanti #610014) on both sides of the PC membrane. Hydrated lipids at a concentration of 1 mM were extruded through the PC membrane 11 times.

### Preparation of supported lipid bilayers

Supported lipid bilayers were formed on 25×75 mm coverglass (IBIDI, #10812). Coverglass was first cleaned with 2% Hellmanex III (Fisher, Cat#14-385-864) heated to 60-70°C in a glass coplin jar and incubated for at least 30 minutes. We washed the coverglass extensively with MilliQ water and then etched with Pirahna solution (1:3, hydrogen peroxide:sulfuric acid) for 10-15 minutes the same day SLBs were formed. Etched coverglass, in water, was rapidly dried with nitrogen gas before adhering to a 6-well sticky-side chamber (IBIDI, Cat# 80608). SLBs were formed by flowing 30 nm SUVs diluted in PBS [pH 7.2] to a total lipid concentration of 0.25 mM. After 30 minutes, IBIDI chambers were washed with 5 mL of PBS [pH 7.2] to remove non-absorbed SUVs. Membrane defects were blocked for 15 minutes with a 1 mg/mL beta casein (Thermo FisherSci, Cat# 37528) diluted in 1x PBS [pH 7.4]. Before use as a blocking protein, frozen 10 mg/mL beta casein stocks were thawed, centrifuged for 30 minutes at 21,370 × g, and 0.22 µm syringe filtered. After blocking SLBs with beta casein, membranes were washed again with 1mL of PBS, followed by 1 mL of kinase buffer before TIRFM.

### Microscopy

For all live-cell imaging experiments, cells were imaged in 1.6 mL of experiment specific imaging media. Base imaging media contained FluoroBrite DMEM (Life Technologies A1896702) supplemented with 25 mM HEPES (pH 7.4) and 1:1000 chemically defined lipid supplement (SF CHIM). Media was then further supplemented with either 10% fetal bovine serum (CHIM) or 0.1% BSA (0.1% BSA CHIM). Alternatively, Ca^2+^ free Ringer’s solution (Ca^2+^ Free) was used, containing 160 mM NaCl, 2.5 mM KCl, 1 mM MgCl_2_, 8 mM glucose and 10 mM NaHEPES, pH 7.5. For treatments, 0.4 mL of experiment specific imaging media containing fivefold final concentration of compound was applied to the dish (or 0.5 ml for a second addition).

Confocal imaging was performed on a Nikon TiE A1R platform with acquisition in resonant mode with a 100x 1.45 NA plan-apochromatic objective. The signal-to-noise ratio was improved by taking 8 or 16 frame averages. Excitation of fluorophores was accomplished using a dual fiber-coupled LUN-V laser launch with 405-nm (BFP), 488-nm (EGFP and NG2), 561-nm (mCherry), and 640-nm (iRFP) lines. Emission was collected on four separate photomultiplier tubes with blue (425-475 nm), green (500-550 nm), yellow/orange (570-620 nm), and far-red (663-737 nm) filters. Blue and yellow/orange channels were recorded concurrently, as were green and far-red. The confocal pinhole was defined as 1.2x the Airy disc size of the longest wave-length channel used in the experiment. In some instances, Nikon Elements denoising software was used to further enhance the signal-to-noise ratio.

For TIRFM and single-molecule imaging (SMol), a separate Nikon TiE platform coupled with a Nikon TIRF illuminator arm and 100x 1.45 NA plan-apochromatic objective was used. Excitation of fluorophores was accomplished using an Oxxius L4C laser launch with 405-nm (BFP), 488-nm (EGFP and NG2), 561-nm (mCherry), and 638-nm (iRFP) lines. Emission was collected through dual-pass filters (Chroma) with blue (420-480 nm) and yellow/orange (570-620 nm) together, and green (505-550 nm) and far-red (650-850 nm) together. An ORCA-Fusion BT sCMOS camera (Hamamatsu) was used to capture images. For TIRFM, images were captured with 2×2 pixel binning. For SMol, the NG2 channel was excited with 100% power for 50 ms from the 488-nm laser in a 16×16 µm region of the PM. Images were registered in rolling shutter mode with 2×2 pixel binning with a 1.5x magnifier lens.

For all types of imaging, Nikon Elements software was used to acquire all images for all experiments and all data was saved with the ND2 file extension.

Membrane binding and lipid phosphorylation reactions reconstituted on supported lipid bilayers (SLBs) were visualized using an inverted Nikon Eclipse Ti2 microscope using a 100x Nikon (1.49 NA) oil immersion TIRF objective. TIRF microscopy images of SLBs were acquired using an iXion Life 897 EMCCD camera (Andor Technology Ltd., UK). Fluorescently labeled proteins were excited with either a 488 nm, 561 nm, or 637 nm diode laser (OBIS laser diode, Coherent Inc. Santa Clara, CA) controlled with a Vortran laser drive with acousto-optic tunable filters (AOTF) control. The power output measured through the objective for single particle imaging was 1-2 mW. Excitation light was passed through the following dichroic filter cubes before illuminating the sample: (1) ZT488/647rpc and (2) ZT561rdc (ET575LP) (Semrock). Fluorescence emission was detected on the iXion Life 897 EMCCD camera position after a Nikon emission filter wheel housing the following emission filters: ET525/50M, ET600/50M, ET700/75M (Semrock). All experiments were performed at room temperature (23°C). Microscope hardware was controlled by Nikon NIS elements.

### Image analysis

Analysis of all images was accomplished using Fiji software[65] using the LOCI BioFormats importer [66]. Custom macros were written to generate channel-specific montages displaying all x,y positions captured in an experiment in concatenated series. In these montages, individual regions of interest (ROIs) were generated around displayed cells.

For confocal images, the ratio of fluorescence intensity between specific compartments was analyzed as described previously [58]. In brief, a custom macro was used to generate a compartment of interest specific binary mask through à trous wavelet decomposition [67]. This mask was applied to measure the fluorescence intensity within the given compartment while normalizing to the mean pixel intensity in the ROI. ROI corresponded to the whole cell (denoted PM/Cell ratio) or a region of cytosol (PM/Cyt), as indicated on the y axis of individual figures.

For TIRFM images, a minimum intensity projection was used to generate ROIs within the smallest footprint of the cells. Background fluorescence was measured and subtracted from all images at all timepoints. The average pixel intensity in each frame (F_t_) was normalized to the mean pixel intensity in the ROI of the time points before treatment (F_pre_) to yield F_t_/F_pre_.

Quantitative data was imported into Prism 8 (GraphPad) for statistical analysis and the generation of graphs and plots. D’Agostino and Pearson normality tests showed data that significantly varied from normal distribution, data were then subjected to a nonparametric Kruskal-Wallis test. If significant difference was found between sample medians, a post hoc Dunn’s multiple comparison test was run.

Representative images were selected based on fluorescence measurements near the median of the sampled population, displayed typical morphology, and robust signal-to-noise ratio. If adjusting brightness or contrast, any changes were made across the entire image.

### Single Molecule Analysis using TrackMate

Mean photon count was estimated using Fiji [65]. Either HEK293A cells expressing PH-PLC81-mNG2×1-3, NG2-PIP4K2A, NG2-PIP4K2B, or NG2-PIP4K2C cells were imaged using SMol settings. Raw images were converted to 32-bit, background subtracted, and grey levels converted into photon counts. These images were then run through Fiji using the TrackMate plugin. Settings for molecule localization were: LoG detector: estimated blob diameter 0.18 µm, threshold 40; initial thresholding by quality; filters on spots: total intensity to match surface localized particles, excluding puncta less than 3; simple LAP tracker: linking max distance 0.5 µm, gap-closing max distance 0.5 µm, gap-closing max frame gap 2. To determine fluorescence intensity per spot, histograms of mean intensity, in each condition, were generated using a 5-photon bin size.

### Kinetic measurements of PI(4,5)P_2_ production

The kinetics of PI(4)P phosphorylation was measured on SLBs formed in IBIDI chambers and visualized using TIRF microscopy as previously described [17]. Reaction buffer contained 20 mM HEPES [pH 7.0], 150 mM NaCl, 1 mM ATP, 5 mM MgCl_2_, 0.5 mM EGTA, 20 mM glucose, 200 µg/mL beta casein (ThermoScientific, Cat# 37528), 20 mM BME, 320 µg/mL glucose oxidase (Serva, #22780.01 Aspergillus niger), 50 µg/mL catalase (Sigma, #C40-100MG Bovine Liver), and 2 mM Trolox (UV treated, see methods below). Perishable reagents (i.e. glucose oxidase, catalase, and Trolox) were added 5-10 minutes before image acquisition. For all experiments, we monitored the change in PI(4)P or PI(4,5)P_2_ membrane density using solution concentrations of 20 nM Alexa647-DrrA(544-647) or 20 nM Alexa488-PLC81, respectively.

### Tables S1 and S2

## Acknowledgments

We thank Robin Irvine for critical reading of the manuscript and valuable suggestions.

## Funding

National Institutes of Health grant 2R35GM119412 (GRVH) National Institutes of Health grant 5F31CA247349-02 (RCW) National Science Foundation CAREER award MCB-2048060 (SDH)

## Author contributions

Conceptualization: RCW, SDH, GRVH

Investigation: RCR, JP, SDH, GRVH

Formal analysis: RCW, JP, SDH, GRVH

Resources: RCR, CPD, JPZ, JP, SDH, GRVH

Funding acquisition: RCW, GRVH

Writing – original draft: RCW, GRVH

Writing – review & editing: RCW, CPD, JPZ, JP, SDH, GRVH

## Competing interests

Authors declare that they have no competing interests.

## Notes

### Competing Interest Statement

The authors have declared no competing interest.

### Summary of Updates

Corrected an error in figure 8b that had arisen during the revision process

